# BIN1 protein isoforms are differentially expressed in astrocytes, neurons, and microglia: neuronal and astrocyte BIN1 implicated in Tau pathology

**DOI:** 10.1101/535682

**Authors:** Mariko Taga, Vladislav A Petyuk, Charles White, Galina Marsh, Yiyi Ma, Hans-Ulrich Klein, Sarah M Connor, Anthony Khairallah, Marta Olah, Julie Schneider, Richard Ransohoff, David A Bennett, Andrea Crotti, Elizabeth M Bradshaw, Philip L De Jager

## Abstract

Identified as an Alzheimer’s disease (AD) susceptibility gene by genome wide-association studies, *BIN1* has 10 isoforms that are expressed in the Central Nervous System (CNS). The distribution of these isoforms in different cell types, as well as their role in AD pathology still remains unclear. Utilizing antibodies targeting specific BIN1 epitopes in human *postmortem* tissue and analyzing RNA expression data from purified microglia, we identified three isoforms expressed specifically in neurons (isoforms 1, 2 and 3) and four isoforms expressed in microglia (isoforms 6, 9, 10 and 12). The abundance of selected peptides, which correspond to groups of BIN1 protein isoforms, was measured in dorsolateral prefrontal cortex, and their relation to neuropathological features of AD was assessed. Peptides contained in exon 7 of BIN1’s N-BAR domain were found to be significantly associated with AD-related traits and, particularly, tau pathology. Since only isoforms 1, 2 and 3 contain exon 7, it appears that decreased protein expression of the N-BAR domain of BIN1 is associated with greater accumulation of tau pathology and subsequent cognitive decline, with astrocytic rather than neuronal BIN1 being the more likely culprit. These effects are independent of the BIN1 AD risk variant, suggesting that targeting specific BIN1 isoforms might be a novel therapeutic approach to prevent the accumulation of tau pathology.

## Introduction

Alzheimer’s disease (AD) is the most common form of aging-related dementia and is characterized by cognitive decline associated with hyperphosphorylation of tau protein, accumulation of amyloid plaques and neurofibrillary tangles, and neuronal loss. In the last few years, several susceptibility loci that contribute to genetic risk for AD, including one that contains the *BIN1* (Bridging integrator 1) gene, have been identified by genome wide association studies (GWAS). The effect size of the *BIN1* variant (tagged by rs6733839) is among the largest for common AD variants; only *APOEε4* and *TREM2*^1 2^ have larger effects. BIN1, a member of the BIN1/amphiphysin/ RVS167 family, is highly expressed in the brain and in skeletal muscle ^3^. It has been implicated in diverse cellular processes such as endocytosis, actin dynamics, DNA repair, membrane trafficking, inflammation and apoptosis ^4^ The *BIN1* gene has 20 exons which encode several known structures including an N-BAR domain, a phosphoinositide (PI) binding motif, a CLAP (clathrin and AP2) binding domain, a Myc-binding domain (MBD) and a Src homology 3 (SH3) domain (Fig. 1a). The N-BAR domain, encoded by exons 1 to 10, is involved in membrane curvature ^5^. The phosphoinositide (PI) binding motif is encoded by exon 11 and is only present in a few BIN1 isoforms (isoform 4, 8 and 12). The CLAP domain is encoded by exons 13 to 16 and is involved in endocytosis ^6^. Its inclusion in mature protein is highly variable among isoforms. Finally, the MBD domain, encoded by exons 17 and 18, plays a role in the regulation of c-Myc, a transcription factor regulating histone acetylation ^7^. *BIN1* is expressed in 14 RNA transcripts, generated by alternative splicing, and 11 of them are translated into respective protein: isoforms 1, 2, 3, 4, 5, 6, 7, 8, 9, 10 and 12 (Fig. 1b). Isoforms 1 through 7 are brain specific with isoform 1 being exclusively expressed in neurons ^3^. Isoform 8 is specifically expressed in muscle ^3^. Isoforms 9 and 10 are ubiquitously expressed and are also detected in the brain ^8^. In the brain, it was reported that BIN1 isoforms are translated into protein in different cell types: neurons mostly express isoform 1, 3, 5 and 7 while astrocytes express isoforms 2, 5, 9 and 10 ^8^.

**Figure 1:**
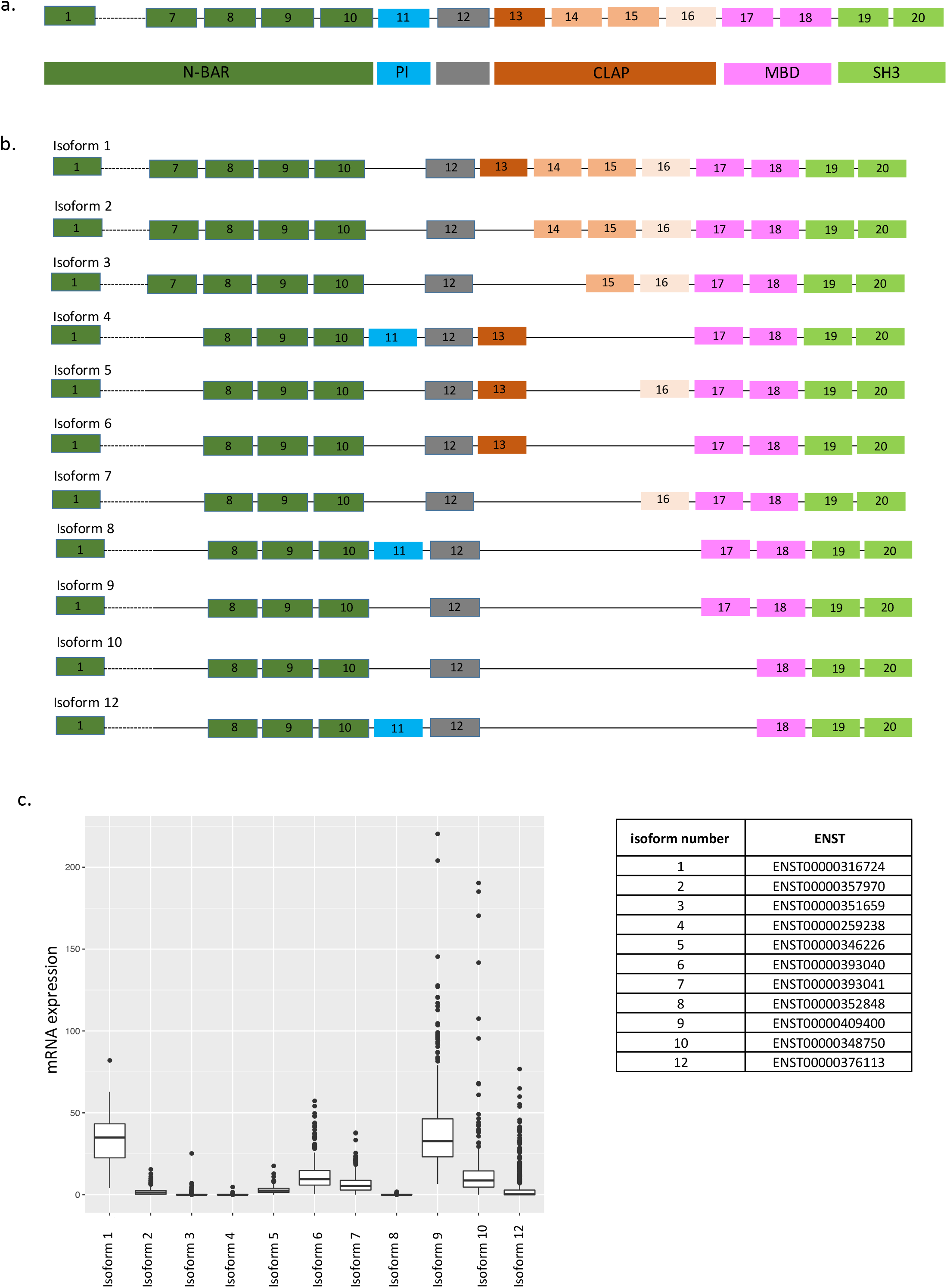
a) BIN1 isoforms schematic representation. b) *BIN1* gene organization and functional domains. c) mRNA expression of BIN1 isoforms in human dorsolateral prefrontal cortex (DLPFC) (n=508)

Genetic analysis of *BIN1* has shown that the rs59335482 insertion allele is associated with an increase of *BIN1* mRNA expression in the brain ^2^, and this manuscript also reports an increase of *BIN1* mRNA expression in the central nervous system (CNS) of AD patients compared to non-AD patients ^2^. These findings highlight that a polymorphism associated with an increase of BIN1 expression in the CNS is not associated with AD risk: the functional consequences of the risk allele remains unclear. A different study found altered methylation in the *BIN1* locus at cg22883290 in relation to neuritic amyloid plaques and a pathologic diagnosis of AD, and this altered methylation pattern was independent of the effect of the rs744373 susceptibility variant ^9^. Thus, there appears to be a convergence of a genetic factor and an environmental or experiential factor mediated by epigenomic changes on risk of AD in the *BIN1* locus. Further, mechanistic dissection has shown that BIN1 interacts with tau ^4^ and is involved in tau-mediated neurotoxicity in a *Drosophila melanogaster*^2^.

In addition to *BIN1*, GWAS have also identified over 30 other loci implicated in AD^10^, and approximately one-third of them are expressed in microglia and other myeloid cells ^10–12^, emphasizing the potential role of immune cells as contributors to the onset of AD. BIN1 has been implicated in inflammation; for example, there is a report of an increase of inflammation in *BIN1*knock-out mice ^13, 14^. However, the role of BIN1 in microglia needs to be clarified as at least some of its isoforms are expressed in all CNS types. In addition, an association between expression of some RNA isoforms and AD was reported: a decrease of isoform 1 and an increase of isoform 9 expression was noted in small collections of human AD *post-mortem* brain samples^15, 16^. De Rossi and colleagues reported that the decrease of isoform 1 (the largest isoform) expression is correlated with a decrease of a neuronal marker, illustrating the loss of neurons observed in AD. On the other hand, the level of isoform 9 seems to be correlated with the abundance of astrocyte (GFAP) and microglia (CD45) markers ^17^. These findings suggest that there may be an increase of isoform 9 expression by reactive astrocytes and microglia during AD. Indeed, BIN1 RNA isoform 9 is highly expressed by glial cells ^18, 19^; however, no study has shown the expression of BIN1 in microglia in *post-mortem* human tissue. Furthermore, the extent of the characterization of the expression of these different isoforms in different cell types had been limited by the small number of commercially available antibodies against exon-selective epitopes of BIN1. Here, using antibodies recognizing specific epitopes of BIN1 and a targeted proteomic approach, we studied the distribution of different translated BIN1 exons and hence protein isoforms in different cell types in the human brain as well as their relation to pathologic measures of AD.

## Material and Methods

### BIN1 isoform cloning

pLVX Ires-Neo lentiviral plasmid from Clontech was cut with EcoRI/MluI to remove the IRES-neo cassette. eGFP-P2A linker expression cassette with a new multiple cloning site was synthesized as linear dsDNA (gBLOCK (IDT)) and cloned into the cut vector using NEB Builder DNA assembly master mix (New England Biolabs). All BIN1 isoforms were then synthesized as gBLOCKs (IDT) and cloned into the GFP-P2A lentiviral vector using NEBuilder DNA assembly master mix according to the manufacturer protocol.

### Cell transfection and Western Blot

HEK293T cells plated in 10-cm dishes were transfected with 12ug of lentiviral vector expressing various BIN1 isoforms using 36ul Lipofectamine 2000 (Life Technologies). After 48 hours, cells were harvested, and protein lysates were prepared from 1×10^6^ 293T cells by adding 100ul of Lysis Buffer (NaCl 140mM, HEPES 10mM pH7.4, NP-40 1%) plus PMSF 1mM, DTT 1mM, Complete Protease Inhibitor Cocktail (Roche). After 1h incubation on ice, lysates were cleared by spinning at 13,000rpm, 10 min, 4°C. Protein concentrations had been determined by BCA protein assay (Biorad). A total of 20μg of protein was separated by SDS-polyacrylamide electrophoresis using NuPAGE 7% Tris-Acetate Gel (Thermo-Fisher), then transferred to a Nitrocellulose membrane. After 1h blocking in 5% (w/v) non-fat milk in TBS with 0.1% Tween 20, the membranes were then incubated overnight at 4°C with in house generated primary antibodies against various epitopes of BIN1 at final concentration of 2ug/ml or anti-BIN1 Ab 1:500 (99D and H100, Santa Cruz), respectively. The membranes were washed in TBST (4×10 min) before 1h incubation with an anti-rabbit HRP 1:5000 (Santa Cruz). The immunoblots were then visualized using the ECL (Amersham).

### Cell culture and TMA preparation

Preparation of cultured cells for paraffin embedding: a batch of 10^8^ 293Expi cells had been transfected with each of the lentiviral vector expressing various BIN1 isoform according to manufacturer’s instruction (Thermo-Fisher). After 48h in culture, cells had been harvested, centrifuged at 1000 rpm at 4°C for 5 minutes, washed in PBS twice. Once PBS had been removed, 10% Neutral Buffered Formalin had been added down the side of the tube onto the cell pellet (Thermo-Fisher). Formalin-Fixed cell pellet had been Paraffin-Embedded (FFPE), sectioned and included Tissue microarray (TMA) blocks in triplicate, ~ 2.25 mm core diameter. After a 24 h fixation period, cells were processed and embedded into paraffin. A tissue microarray (TMA) was constructed from formalin-fixed, paraffin-embedded cell pellet preparations. Triplicate 2.25 mm cores from each of the cell pellet blocks were punched and placed into the TMA block. Untransfected 293 cell pellet preparatiom was also included in triplicate. A non-related tissue sample was included in the TMA as an orientation marker.

### Validation of isoform-specific anti-BIN1 antibodies

Five-micron thick sections of TMA block containing 11 cell pellets were prepared and placed on charged slides. Immunohistochemical staining was performed using Ventana Discovery Ultra automated staining platform (Roche Ventana Medical Systems, Tucson, AZ). Briefly, paraffin sections deparaffinized and rehydrated. Epitope retrieval was performed in Ventana CC1 (EDTA, pH 8.0) buffer at 95 °C for 64 min. Slides were reacted against anti-BIN1 antibodies at one concentration (final concentration of 0.25-1 μg/mL, as selected from immunohistochemistry method optimization studies). Following primary antibody incubation, sections were incubated with secondary goat anti-rabbit antibodies (OmniMap HRP kit, Roche Ventana) followed by detection with diaminobenzidine (DAB) and counterstaining with hematoxylin. Stained slides were digitized, and staining was evaluated visually. Photomicrographs were captured at 100x magnification using ImageScope software (Leica Microsystems). A non-related tissue sample was included as an orientation marker.

### ROSMAP cohort

The Religious Orders Study (ROS) and Memory and Aging Project (MAP) are both cohort studies directed by Dr. David Bennett at RUSH University. Non-demented individuals over age 65 are recruited and consent to brain donation at the time of death. Both cohorts implement the same annual evaluations including 19 cognitive functional tests and use accepted and validated procedures to diagnose AD and other dementias and mild cognitive impairment (MCI). Since 1994 (ROS) and 1997 (MAP), more than 3,069 individuals have been enrolled and 1641 are still alive and the follow-up rate exceeds 95% and the autopsy rate exceeds 90%. All participants undergo a uniform structured evaluation for AD including CERAD, Braak Stage, NIA-Reagan, and a global measure of AD pathologic burden with modified Bielschowsky; amyloid load and PHFtau tangles by immunocytochemistry and image analysis and stereology; Lewy bodies and TDP-43 by immunocytochemistry, and the age, location, and size of all macro- and microscopic infarcts. Both frozen and fixed brain tissue is available on request from these subjects. Our laboratory has generated DNA methylation scans, H3K9Ac ChipSeq, and RNAseq data (n=508) from the frozen dorsolateral prefrontal cortex as part of other studies (Suppl. Table 1). We also have RNA-seq data from FACS-purified viable microglia from some of these individuals at autopsy (n=21).

### Immunohistochemistry on human *post-mortem* brain tissue

Formalin-fixed *post-mortem* brain tissues were obtained from brains donated to Rush University Medical Center. All participants consented to brain donation at the time of death. Six μm sections of formalin-fixed paraffin-embedded tissue from the cortex frontal (n=3) were used to stain NeuN (Millipore), IBA1 (Wako), ALDH1L1(eBioscience), CD45 (Novus Biological), along with anti-BIN1 antibodies provided by Biogen.

Immunohistochemistry was performed using citrate for antigen retrieval. The sections are blocked with blocking medium (8% of horse serum and 3% of BSA) and incubated overnight at 4°C with primary antibodies. Sections are washed with PBS and incubated with fluorochrome conjugated secondary antibodies (Thermo Fisher) and coverslipped with anti-fading reagent with Dapi (P36931, Life technology). Photomicrographs are captured at X20 magnification using Zeiss Axio Observer.Z1 fluorescence microscope and exported to Image J imaging software (NIH, Maryland, USA). The images have been quantified using CellProfiler and CellProfiler Analyst.

### Isolation of human monocytes and MDMi cell differentiation

Peripheral blood mononuclear cells (PBMC) were extracted from blood samples from the PhenoGenetic cohort using a standard Ficoll protocol (Ficoll-PaqueTM PLUS). PBMCs were frozen at a concentration of 1-3×10^7^ cells/mL in 10%DMSO + 90%FBS and stored in −80°C until they are processed. CD14+ monocytes are purified from frozen PBMC samples using a positive isolation strategy (CD14+ selection kit, Miltenyi). 100,000 monocytes were allocated per well in 96-well plates. To differentiate to MDMi, monocytes were cultured in RPMI-1640 media supplemented with 10ng/ml M-CSF, GM-CSF and NGF-β, as well as 100ng/ml CCL2 and IL-34 for 10 days.

### Expression Analysis by Real-Time RT-PCR

Total RNA was isolated using the RNeasy Plus Micro Kit (Qiagen, Germany), according to the manufacturer’s protocol. The expression of genes of interest were measured by qRT-PCR, in duplicate. B2M was used as an internal control. 5μL of the cDNA was amplified using 10μL of TaqMan Fast Advanced Master Mix, 1.5 of primers and 4μL of RNase free water. qRT-PCR was run on Quantstudio 3 detection system (Applied Biosystems, USA).

### Isolation of adult human microglia, RNA-sequencing and data processing

Microglia were isolated from autopsy brain material as described in Olah *et al.*, Nature Communication, 2018. Briefly, upon arrival of the autopsy brain sample, the cerebral cortex and the underlying white matter were dissected under a stereomicroscope. All procedures were performed on ice. Only microglia isolated from the grey matter of the dorsolateral prefrontal cortex (DLPFC) were used in this study (n=21). The dissected tissue was first mechanically dissociated. Subsequently, myelin was depleted by the use of anti-myelin magnetic beads, following which the cell suspension was enriched in microglia with anti-CD11b magnetic beads. Microglia were then sorted based on their characteristic CD11b and CD45 expression and a viability dye (7AAD) on a BD FACS Aria II sorter. Cells were sorted into a 96-well PCR plate containing lysis buffer. Following FACS the lysate was snap frozen on dry ice and stored at −80°C until further processing. Library construction was performed using the SmartSeq-2 protocol described in detail elsewhere (Olah et al.). Samples were sequenced on an Illumina HiSeq2500 platform with a read length of 100bp and paired end reads. RNA-Seq reads in FASTQ format were inspected using FASTQC program. Barcode and adapter contamination, low quality regions (10bp at beginning of each fastq reads) were trimmed using FASTX-toolkit. The STAR (v2.5.3a) aligner software was used to map paired-end reads to the reference genome (assembly GRCh38) using Ensembl gene annotation (release 91). Transcription levels were estimated using the RSEM (v1.2.31) software and reported in Transcripts Per Million (TPM). Four out of 25 samples were removed because of low percentage of exonic and UTR reads (< 30%) or low total number of reads mapping to the transcriptome (< 106 reads).

### SRM proteomics

Selected Reaction Monitoring (SRM) proteomics was performed using frozen tissue from dorsolateral prefrontal cortex (DLPFC) (Suppl. Table 1) (n=1377). The sample preparation for LC-SRM analysis follows standard protocol, as described elsewhere ^20, 21^. An average ~20 mg of brain tissue from each subject was homogenized in denaturation buffer (8M urea, 50 mM Tris-HCl pH 7.5, 10 mM DTT, 1 mM EDTA). Following denaturation, 400 ug protein aliquots were taken for further alkylation with iodoacetamide and digestion with trypsin. The digests were cleaned using solid phase extraction, following readjustment of tryptic peptide digests concentration to 1 ug/uL. 30 uL aliquots were mixed with 30 uL synthetic peptide mix. All liquid handling steps were performed in 96-well plate format using Epmotion 5075 TMX (Eppendorf) and Liquidator96 (Rainin). We selected 7 proteotypic peptides based on BIN1 isoform-specificity and peptide detectability prediction tool CONSeQuence ^22^ (Suppl. Table 2). The 7 synthetic heavy peptides labeled with ^13^C/^15^N on C-terminal lysine and arginine were purchased from New England Peptide (Gardner, MA) with cysteine modification with carbamidomethylation. All LC-SRM experiments were performed on a nano ACQUITY UPLC coupled to TSQ Vantage MS instrument, with 2 μL of sample injection for each measurement. A 0.1% FA in water and 0.1% in 90% ACN were used as buffer A and B, respectively. Peptide separations were performed by an ACQUITY UPLC BEH 1.7 μm C18 column (75 μm i.d. × 25 cm) at a flow rate 350 nL/min using gradient of 0.5% of buffer B in 0-14.5 min, 0.5-15% B in 14.5-15.0 min, 15-40% B in 15-30 min and 45-90% B in 30-32 min. The heated capillary temperature and spray voltage was set at 350 °C and 2.4 kV, respectively. Both the Q1 and Q3 were set as 0.7 FWHM. The scan width of 0.002 m/z and a dwell time of 10 ms were used. All the SRM data were analyzed by Skyline software ^23^. All the data were manually inspected to ensure correct peak assignment and peak boundaries. The peak area ratios of endogenous light peptides and their heavy isotope-labeled internal standards (i.e., L/H peak area ratios) were then automatically calculated by the Skyline software and the best transition without matrix interference was used for accurate quantification. The peptide relative abundances were log2 transformed and centered at the median.

## Results

### Study of the distribution of BIN1 isoforms in the CNS using transcriptomic and proteomic approaches

We first evaluated the distribution of BIN1 isoforms in the CNS by analyzing RNA expression of each isoform in human dorsolateral prefrontal cortex (DLPFC) (Fig. 1c); the figure also provides the reference numbers for each BIN1 isoform and an outline of the locus. Isoforms 1 and 9 are most highly expressed, followed by isoforms 6, 7 and 10. Isoforms 2, 3, 5, and 12 are expressed at low levels, and isoform 8, which has been reported to be specific to muscle ^3^, is absent.

To characterize the cell-specific expression of BIN1’s different isoforms at the protein level in human *post mortem* tissue, 4 antibodies against 4 specific exons (exon 7, 11, 13 and 16) have been generated. The targeted exons are present in different domains of BIN1: exon 7 is present in the N-BAR domain; exon 11 is in the PI domain; while exons 13 and 16 are in the CLAP domain (Suppl. Fig. 1a). These exons are alternatively spliced in selected isoforms (Fig. 1b). To validate the various BIN1 antibodies, we cloned BIN1’s 11 protein-coding isoforms and expressed them in HEK293T cells. As positive controls, we used commercially available anti-BIN1 antibodies: clone 99D (recognizing a mid-portion of BIN1 present in isoforms 1 through 9) and clone H100 (recognizing the C-terminus of BIN1 present in isoforms 1 through 12). No basal expression of BIN1 was detected in the HEK293T cell clone that we used to express the different isoforms (data not shown). As expected, the 4 custom antibodies against different epitopes of BIN1 recognized the selected isoform, according to the scheme depicted in Fig 1b (Suppl. Fig. 1b). Furthermore, the 4 custom antibodies against different epitopes of BIN1 recognized the selected isoform expressed in 293T cells by immunohistochemistry (Suppl. Fig. 2). Specific reactivity was demonstrated against the appropriate transfected cell pellet controls, while no non-specific reactivity was observed (Suppl. Fig. 2). Having validated these new reagents, we deployed them to stain human *post-mortem* brain samples to characterize the distribution of the different isoforms of BIN1 by immunofluorescence.

### BIN1 isoforms are expressed in neurons and astrocytes

Immunostaining using the antibody recognizing exon 7 showed a co-localization with NeuN, a neuronal marker and with ALDH1L1, an astrocyte marker (Fig. 2a). This observation suggests that exon 7 of BIN1 is expressed in both neurons and astrocytes. The same approach was used for the antibodies recognizing exons 11, 13 and 16 in order to define the cell types expressing these exons. Exons 11, 13 and 16 are expressed in both neurons and astrocytes (Fig. 2b, 2c, 2d); however, only a subset of astrocytes expressed exon 16 of BIN1 (Fig. 2d, arrows), suggesting a more context-specific role for this isoform. Considering that exon 7 is only present in BIN1 isoforms 1, 2, and 3 (Fig. 1b), the results indicate that these three isoforms might be expressed in neurons and astrocytes. BIN1 isoforms 4 and/or 12, which both contain exon 11 (Fig. 1b), are expressed in neurons and astrocytes. Similarly, isoforms 4, 5 and 6 that contain exon 13 and isoform 7 which contains exon 16 (Fig. 1b) appear to be expressed in both cell types. Since isoform 8 is not expressed in the CNS (Fig. 1c), we excluded it from this annotation relating to neurons and astrocytes.

**Figure 2:**
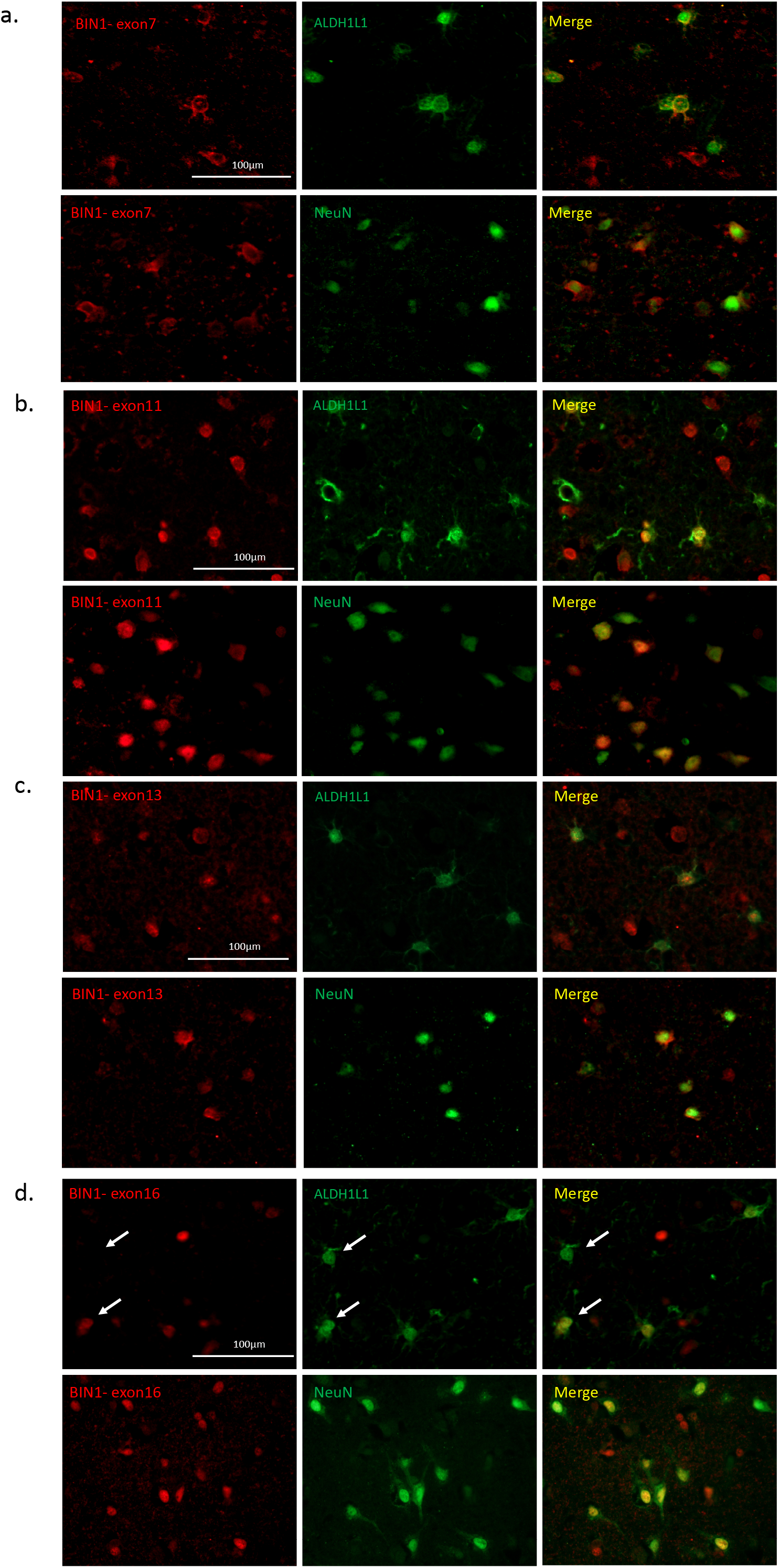
Characterization of different BIN1 isoform pattern of expression in different brain cells. a-d) Co-immunostaining using antibodies recognizing specific exons of BIN1 (exons 7, 11, 13 and 16) in red with neuronal (NeuN) and astrocyte (ALDH1L1) markers in green in human *post-mortem* brain tissue. e) Summary of the distribution of BIN1 exons in different cell type in human *post-mortem* brain tissue (n=3).

### BIN1 isoforms 6, 9, 10 and 12 are expressed in human microglia

Two previous publications have tried to characterize the presence of BIN1 in *post-mortem* human brain samples, but neither had shown expression in microglia ^17, 18^. In order to investigate whether BIN1 is expressed in microglia, we performed a BIN1 mRNA isoform expression analysis in transcriptomes of 21 subjects with live grey matter microglia purified from fresh autopsy ^11^ (Fig. 3a). Using estimates of isoform abundance derived from our paired-end reads of 100 bp, we observed expression of isoforms 6, 9, 10 and 12 in these purified cortical microglia, with RNA isoform 9 being the most highly expressed isoform in both batches of samples. Estimates of isoform expression level from these RNA sequencing data are supported by the results of an alternative approach which counts the number of reads mapped to each exon (Suppl. Fig. 3a). Overall, these results support the presence of BIN1 RNA in purified human microglia, and shotgun proteomic data generated from purified microglia derived from other human samples by the same pipeline confirmed the detection of BIN1 at the protein level. As seen in Suppl. Table 3, 6 BIN1 peptides encoded by different exons were present in these data.

**Figure 3:**
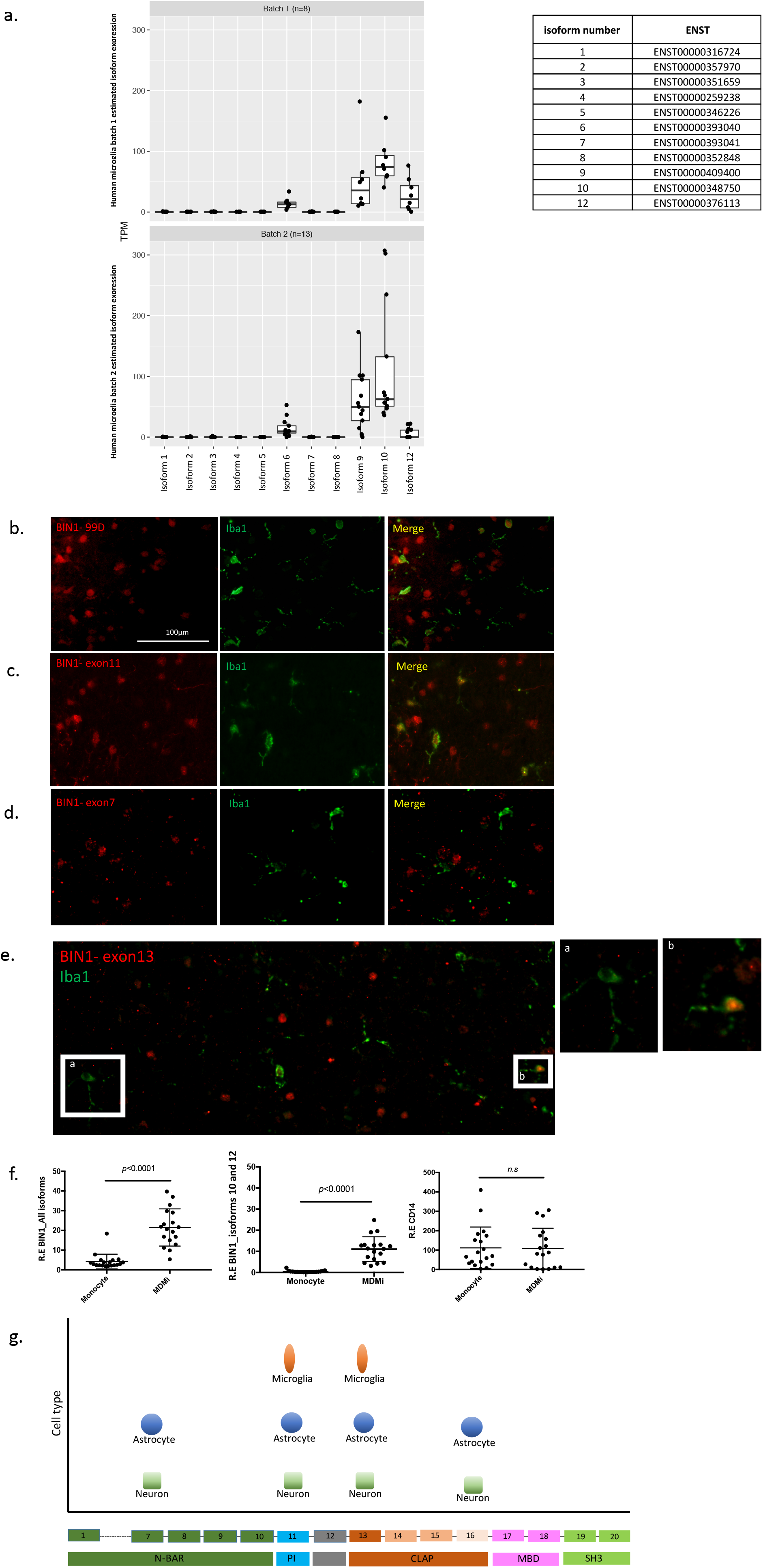
a) RNA expression of specific BIN1 isoform expressed in microglia from human fresh brain tissues. Two different batches of microglia preparation showed comparable results (n=21). b-e) Co-immunostaining of antibodies recognizing specific exons (exons 7, 11 and 13) (red) and BIN1 99 recognizing isoforms 1-9 (red) with Iba1, a microglia marker (green) in *post mortem* human brain tissue (n=3). f) Analysis of gene expression level of BIN1 (all isoforms and isoforms 10 and 12) in monocytes and MDMi (n=19). No difference of gene expression level has been reported for CD14 between two cellular models. Relative gene expression is reported in Y axis. g) Summary graph illustrating the distribution of BIN1 exons in different cell types in CNS.

In order to explore the expression of BIN1 by microglia in human *post mortem* brain tissue, an antibody recognizing an epitope of BIN1 contained within isoforms 1-9 (Ab BIN1 99D) was used in combination with the microglial marker IBA1 (Fig. 3b). The results showed expression of BIN1 in the nucleus of IBA1 positive cells, suggesting the presence of BIN1 isoforms 1-9 in microglia. In order to narrow down the identity of the isoforms that are specifically expressed by microglia at the protein level, the exon specific antibodies of BIN1 were again utilized. The BIN1 antibody recognizing exon 11, contained in isoforms 4, 8 and 12 (Fig. 3c), produced a positive staining in the nucleus of IBA1 positive cells, confirming the expression of isoform 12 which was also observed at the mRNA level. Isoforms 4 and 8 also contain exon 11, but they can be excluded since we didn’t detect these isoforms at the RNA level in purified microglia (Fig. 3a). Subsequently, we tested a BIN1 antibody against exon 7 (which is present in isoforms 1, 2 and 3) but observed no co-localization between BIN1 and IBA1 (Fig. 3d), suggesting that BIN1 isoforms 1, 2 and 3 are not expressed by microglia. These results confirm our mRNA expression analysis of microglia where none of these isoforms were detected (Fig. 3a) and are consistent with previously published reports of isoform 1 and 3 expression in neurons and isoform 2 in astrocytes ^8^. Finally, we tested an antibody against exon 13 (present in isoforms 4, 5 and 6). As observed in Fig. 3e, we detected positive staining in the nucleus of a few IBA1 positive cells, suggesting that only a subset of microglia express BIN1 containing exon 13. These results confirm our RNA expression analysis of microglia, where isoform 6 but not isoforms 4 and 5 was detected. In summary, using antibodies against selected BIN1 epitopes, we find that BIN1 isoforms 6, and potentially 9 and 12 are expressed by microglia in the aged human brain.

To explore whether these isoforms are expressed in all myeloid cells (including microglia and monocytes) or more specific to myeloid cells of the CNS, we evaluated data from human purified monocytes and matched Monocyte Derived Microglia-like (MDMi) cells (Fig. 3f). We note a significant increase of BIN1 isoforms expressed in microglia as isoforms 10 and 12 (p<0.0001) in MDMi compared to their corresponding monocytes. These findings showed that isoforms present in myeloid cells are expressed in a context-dependent manner and are probably more prominent in microglia than in peripheral myeloid cells. Fig. 3g presents a summary of these results.

We evaluated the possible role of genetic variants on BIN1 expression; in particular, we evaluated the BIN1 single nucleotide polymorphisms (SNPs) previously reported to be associated with Alzheimer’s Disease (AD). However, these variants had no effect on gene or isoform mRNA expression in a well-powered sample of cortical tissue data ^24^ (n=503) or in a small sample of purified microglia (n=20) (Suppl. Fig. 3b and 3c). Other non-AD SNPs in the BIN1 locus were seen to have a modest effect on expression in cortical tissue data for isoform 10, which is expressed in microglia (Suppl. Tables 4a and 4b), but this was not seen in the small dataset derived from purified microglia.

### Association between BIN1 isoforms and AD-related traits

Using RNA sequencing data from the dorsolateral prefrontal cortex (DLPFC) of 541 participants in two prospective studies of aging (ROSMAP subjects), we assessed the relation of BIN1 isoforms to AD-related traits (Suppl. Fig. 4). We find no significant association between *BIN1* isoforms with a syndromic diagnosis of AD dementia and AD-related traits such as a person’s slope of cognitive decline prior to death or quantitative measures of neuritic amyloid plaques, neurofibrillary tangles or PHF tau (which measures the presence of an AD-associated phosphorylated epitope) burden.

Given potential differences between RNA and protein expression, we then turned to a quantitative proteomic approach: we used signal reaction monitoring in DLPFC samples of a larger set of ROSMAP participants (n=1377) to measure the abundance of 7 peptides that originate from different domains of the BIN1 protein (Fig. 4 and Suppl. Tables. 2-3). Five of the peptides map to the N-BAR domain: two in exon 7, one spanning exon 7 and 8 and one each in exons 8 and 10. The other two peptides map to a linker domain (encoded by exon 12) and exon 13 which contains the proximal element of the CLAP domain.

**Figure 4:**
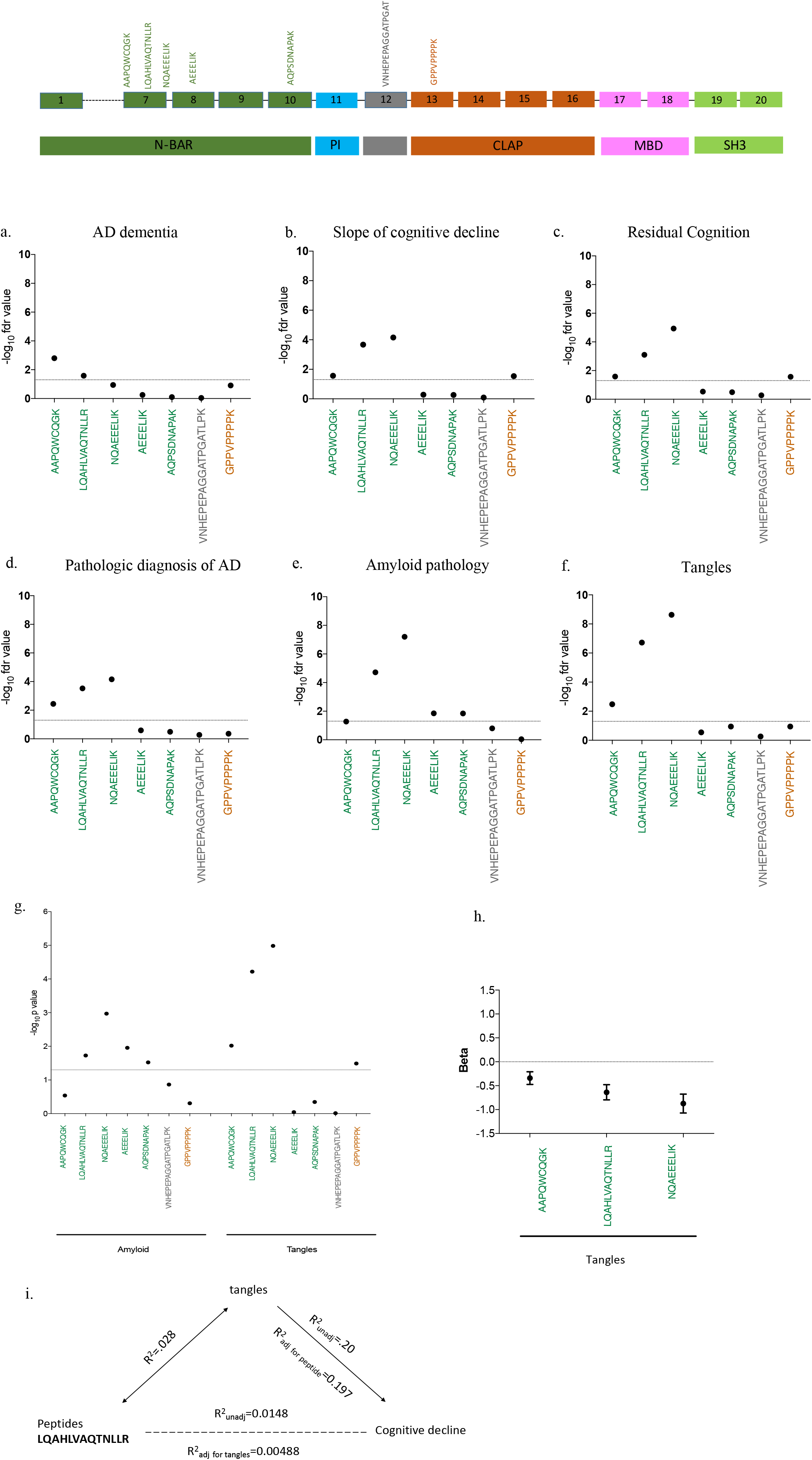
a-f) The abundance of each of 7 peptides from 3 different domains of BIN1 have been associated with cognition and AD pathology in human DLPFC by quantitative proteomic approach. g) Association between the abundance of BIN1 peptides and AD pathology after adjusting for tangles and amyloid pathology. h) Correlation between BIN1 peptides localized in exon 7 and tangles. i) Graph illustrating the association between BIN1 peptides (i.e LQAHLVAQTNLLR) and cognitive decline is not independent of tangles.

After testing 7 peptides (Suppl. Fig 5), only the three peptides which contain sequence from exon 7 display associations with AD related-traits: a diagnosis of AD dementia, the participants’ trajectory of cognitive decline, residual cognition and quantitative measures of both amyloid and Tau pathology (tangles) (Fig. 4a-f and Suppl. Table 5). In secondary analyses, we also evaluated other neuropathologic traits that are available in these subjects and find no association with neurovascular measures, cerebral amyloid angiopathy, hippocampal sclerosis, Lewy Bodies, or the burden of TDP43 pathology (Suppl. Table 6). These findings suggest a specific association between exon 7 of BIN1 and AD-related pathologies, and our measures of amyloid and tau pathology appear to be more strongly associated than the measures related to clinical function. Overall, as pathology accumulates, exon 7 peptide expression is reduced. Similarly, the expression of these peptides is diminished as cognitive functions worsens. To address the possibility that these associations are due to a shift in cortical cell populations, we repeated these analyses in a subset of individual in which we could estimate the proportion of different cell types using RNA sequence data (n=508). As seen in Supplementary Table 7, the results of these analyses remain significant for AD pathologies after including a correction for the different cell types.

Since the burden of amyloid and tangles are partially correlated to one another, we used conditional analyses to assess whether BIN1 exon 7 peptides are primarily driven by one of the two pathologies or are independently associated to both measures (Suppl. Table 8). When we adjust for the burden of tangles, most of the association with amyloid is no longer significant, aside for the peptide that bridges exons 7 and 8, which is diminished by 49.2% (Fig. 4g). On the other hand, the associations with tangles are diminished but remain significant when we account for the effect of amyloid, suggesting that the association of the alternatively spliced exon 7 of BIN1 with AD is primarily driven by tangles, with a possible minor component involving amyloid independent of tau (Fig. 4g).

Since the primary effect of BIN1 exon 7 on pathology is exerted through tangles, we further explored whether this relation with tangles explained the association of BIN1 with cognitive decline; in evaluating different models using mediation analyses, we find that the peptide LQAHLVAQTNLLR explains 1.48% of the variance in cognitive decline (*p*=4.63.10^−5^), when modeled independently: with higher protein expression of this peptide we see slower cognitive decline. On the other hand, tangles explain 20% of the variance in cognitive decline (*p*=7.59.10^−67^), where higher tangles increase the rate of cognitive decline. When modeled jointly, the effect of the peptide LQAHLVAQTNLLR on cognitive decline is reduced significantly (r^2^ from 1.48% to 0.49%), whereas the effect of tangles on cognitive decline is unchanged (r^2^ from 20% to 19.7%), suggesting that the majority of the role of BIN1 in cognitive decline is not independent of Tau (Suppl. Table 9). In other words, we propose that reduced inclusion of exon 7 into BIN1 proteins contributes to increases in tangle burden (Fig. 4h) which then causes more rapid cognitive declines. Fig. 4i presents a summary of these results.

### Effect of genetic variants and known risk-associated molecular features on peptide measures

We evaluated the role of *APOEε4* which has a strong role in AD, with both tangles and cognitive decline: specifically, *APOEε4* explains approximately 7.7% of the variance in tangles while the BIN1 peptide LQAHLVAQTNLLR explains up to 2.8% of the variance in tangles. In an independent model containing the peptide, LQAHLVAQTNLLR peptide explains 1.48% of the variance in cognitive decline, while APOEε4 explains 5.7%. However, in a conditional model containing both LQAHLVAQTNLLR peptide and APOEε4, the LQAHLVAQTNLLR peptide explains 0.76% of the variance in cognitive decline (Fig. 5b), whereas, APOEε4 explains 6% of the variance in cognitive decline. This indicates that the association of LQAHLVAQTNLLR peptide with cognitive decline persists after adjustment with APOEε4 (p=0.00386) but is not entirely independent.

**Figure 5:**
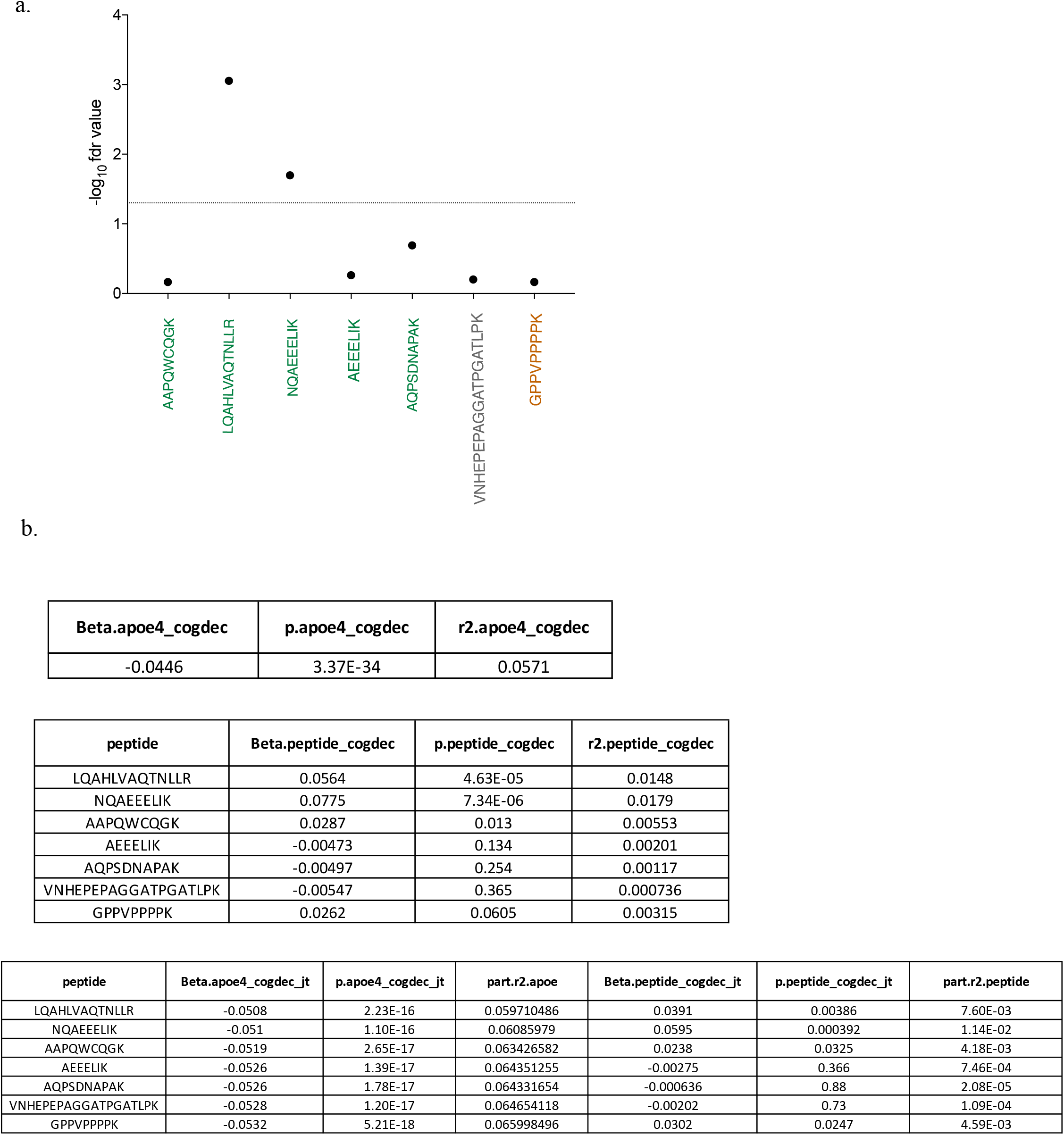
a) BIN1 peptides LQAHLVAQTNLLR and NQAEEELIK from exon 7 are associated with APOE*ε4*. b) Correlation between BIN1 peptides and cognitive decline after adjusting with APOE*ε4* showing that the relationship between BIN1 peptides and cognitive decline is not independent of APOE*ε4*.

Secondarily, we evaluated whether the other known AD susceptibility variants influence BIN1 peptides, but we found no evidence of association between AD variants and BIN1 peptides after correcting for the testing of multiple hypotheses (Suppl. Table 10). Further, we performed a cis-pepQTL (+-1Mb) (peptide QTL) analysis of the BIN1 locus but found minimal effects of genetic variation in the vicinity of BIN1 on peptide expression. We found a single linkage disequilibrium (LD) block where the top SNP rs74490912 is associated with expression of BIN1 peptide LQAHLVAQTNLLR (p=4.9.10-6); but this SNP is in only weak LD with the BIN1 AD SNP rs6733839 (R^2^=0.156).

In other analyses, we have found Tau pathology to be associated with certain miRNA (miR132 and mir129) (https://doi.org/10.1101/234351) as well as certain modules of co-expressed genes derived from cortical RNA sequence data ^24^.We therefore extended our analyses to include these variables and find that the association of the BIN1 peptide LQAHLVAQTNLLR with tangles is independent of all of these factors (Suppl. Table 11). Finally, we have recently reported large-scale changes in the neuronal epigenome in relation to Tau pathology ^25^, and we have developed a person-specific score for the extent of cortical epigenomic alteration. Using this score, we found that there is no association between tau-related epigenomic alterations, as defined by the H3K9Ac histone marks, and the LQAHLVAQTNLLR or other BIN1 peptides (Suppl. Table 12). Thus, the relation of BIN1 protein expression to the accumulation of Tau pathology appears to be independent of these factors.

### Validating the association of Exon 7 with AD in human *post-mortem* tissue

Given the association of exon 7 with AD and AD-related pathologies in the proteomic analyses, we stained AD and cognitively non-impaired subjects with the antibody recognizing BIN1 exon 7 and ALDH1L1, a marker of cortical astrocytes, or NeuN, a neuronal marker to resolve which cell type may be driving the association. We collected images in a systematic manner and used Cell Profiler to automatically segment these images and count the number of cells expressing each marker combination. We then compared the frequency of cells expressing each marker combination in the AD and cognitively non-impaired subjects (Supplementary Figure 6). While the number of BIN1 exon 7+ neurons shows no difference, we find a significant decrease in the number of astrocytes expressing BIN1 exon 7 in AD compared to cognitively non-impaired subjects while there is not a significant difference in the total number of astrocytes between the two classes of subjects. This suggests that the role of BIN1 exon 7 in AD is related to its role in astrocytes and perhaps that loss of a class of BIN1 exon 7+ astrocytes contributes to the accumulation of Tau pathology.

## Discussion

Since its identification as a genetic risk factor for AD in 2010, several studies have focused on characterizing the mechanism by which BIN1 contributes to the onset of AD ^2, 15, 26^. Currently, only 2 of the 11 translated isoforms have been described as being differentially expressed in the AD brain: a decrease of isoform 1 and an increase of isoform 9 expression correlate with AD ^15^. However, it remains unclear (1) whether these transcriptional changes are due to differences in cell type proportions and (2) in which cell type(s) BIN1 may play a role in influencing AD risk since it is widely expressed.

In this study, we identified the cell types expressing different BIN1 RNA and protein isoforms and investigated their association with AD neuropathology. At the RNA level, we detected expression of isoforms 1, 2, 3, 4, 5, 6, 7, 9, 10 and 12 in the human DLPFC, and we observed that isoforms 1 and 9 are highly expressed compared to the other isoforms. Utilizing antibodies specifically targeting exon 7 on *post-mortem* human brain, we deduced that isoforms 1, 2 and 3 are expressed in both neurons and astrocytes. Exon 7 encodes the N-BAR domain of BIN1 and is specific to isoforms 1, 2 and 3 ^5, 27^; Exon 7 enables the interaction between BIN1 and dynamin 2 (DNM2), a ubiquitously expressed GTP binding protein, and this interaction facilitates endocytic uptake ^28^. Absence of exon 7 leads to the inhibition of uptake, suggesting that the BIN1 N-BAR domain plays a key role in endocytosis via its interaction with DNM2 ^28^. In addition, a decrease of DNM2 mRNA in AD temporal cortex has been reported ^29^. In our study, we have shown that the expression level of exon 7 peptides (AAPQWCQGK, LQAHLVAQTNLLR, NQAEEELIK) are associated with tau pathology and cognitive decline. In addition, we report that exon 7 peptides of BIN1 are negatively correlated with tangles, suggesting that decreased expression of neuronal/astrocyte isoforms 1, 2, and 3 contributes to greater accumulation of tangles and cognitive decline. Since we found no association between BIN1 isoforms and AD traits at the RNA level, it appears that the alteration in BIN1 protein isoforms may be occurring post-translationally. Interestingly, BIN1 has been reported to regulate endocytosis in neurons, and the loss of its function appears to promote the propagation of tau pathology in an *in vitro* model, supporting our findings ^30^. However, our *in situ* validation revealed dysregulation of expression of BIN1 exon 7 in astrocytes in AD subjects suggesting that the lack of expression of these specific BIN1 isoforms in astrocytes might be involved in AD pathology.

In summary, based on our data and evidence from the literature, we propose that decreased expression of isoforms 1, 2, and 3 yields increases in the endocytosis process and promotes the formation the phosphorylation of AD-related epitopes in neurons, either in a cell autonomous or non-cell autonomous fashion. The potential function of these isoforms expressed in astrocytes should be investigated further. This model implicates BIN1 in the narrative of altered endocytosis that is suggested by several of the known AD susceptibility loci. However, the BIN1 AD variant itself does not appear to function through these alterations in the splicing of the N-BAR domain: the rs6733839 variant’s mechanism remains unclear and independent of the role of the N-BAR domain in AD.

Given our results, BIN1’s expression in microglia may be unrelated to the convergence of other AD risk factors in microglia, although we need larger microglial-specific datasets to fully explore the function of microglial BIN1 in AD. Nonetheless, we have enriched our understanding of BIN1 in this study: our multidisciplinary approach identified 4 specific BIN1 protein isoforms (isoforms 6, 9, 10 and 12) expressed by these cells. Except for isoform 6, these isoforms are characterized by the lack of a CLAP domain, and they appear to be expressed at higher level in microglia-like cells than in peripheral myeloid cells. The CLAP domain has been described to be involved in clathrin mediated endocytosis and in the recruitment of dynamin. The absence of exon 7 and the CLAP domain in BIN1 isoforms expressed by microglia suggest that BIN1 may have a function during aging and AD that is unrelated to clathrin mediated endocytosis. BIN1 has been implicated in phagocytosis in macrophages ^13^, and an increase of incidence of inflammation has been reported in BIN1 knock out mice ^14^, suggesting a possible anti-inflammatory function of BIN1. In addition, we show that BIN1 isoforms 10 and 12 are expressed in monocyte-derived microglia-like cells and not in peripheral monocytes, highlighting the existence of context-specific expression of BIN1 protein isoforms by microglia.

Overall, using multiple approaches, our study has refined the cell-type specific expression patterns of the different BIN1 isoforms at the protein level. Our findings showed a strong association between BIN1 isoforms expressed by neurons/astrocytes and tangles that contributes to cognitive decline in AD, and our *in situ* studies refined these association to prioritize astrocytes as the target cell type. While these isoforms may be unrelated to the effect of the rs6733839 AD risk variant, they nonetheless represent a distinct contribution of this complex protein to AD, offering new insights into how BIN1 could be targeted or could serve as an outcome measure in AD studies.

### Ethical approval

“All procedures performed in studies involving human participants were in accordance with the ethical standards of the institutional and/or national research committee and with the 1964 Helsinki declaration and its later amendments or comparable ethical standards.”

## Acknowledgements

Biogen Translational Pathology Laboratory members Michael Craft, Shanqin Xu, Stefan Hamann, provided technical assistance with the histopathological work and image analysis. This work was supported by a sponsored research agreement from Biogen to Dr. De Jager.

## Authors Contributions

P.D.L, E.M.B, R.R, A.C and M.T conceived the study and wrote the manuscript. A.C and M.T designed and validated the antibodies. G.M validated the antibodies by immunohistochemistry. V.A.P. performed and analyzed the Selected Reaction Monitoring proteomics data. C.W performed the biostatistical analysis. M.O generated the purified human microglia data which was analyzed by H.U-K and Y.M. J.S and D.B provided human brain sample. S.M.C and A.K provided technical help.

## Competing Financial Interests

A.C. and G.M. are employees of Biogen. R.M.R. is former of Biogen employee. R.M.R. is an employee of Third Rock Venture Capital.

## Supplementary figures

**Suppl. Figure 1:** a) Schematic illustration of BIN1 antibodies epitopes specificity, recognizing exons 7, 11, 13 and 16. b) Western Blot showing 4 custom antibodies and 2 commercially available antibodies targeting different epitopes of BIN1 and recognizing its isoforms in HEK293T cell clones expressing each BIN1 isoform respectively.

**Suppl. Figure 2:** Validation of isoform-specific anti-BIN1 antibodies using BIN1 Isoform 1-12 transfected cell pellets. Representative images of IHC staining of formalin-fixed, paraffin embedded 293 cells transfected with BIN1 isoforms 1-12 (panels #1-12), or non-transfected 293 cells (panel N).

**Suppl. Figure 3:** Illustration of the number of reads mapped to each BIN1 exon in human microglia from fresh DLPFC tissue. The number of read mapped to the exon is reported on Y axis.

**Suppl. Figure 4:** BIN1 isoforms have been associated with cognitive slope, amyloid pathology, neuritic plaques neurofibrillary tangles and tangles at RNA level.

**Suppl. Figure 5:** Abundance of 7 BIN1 peptides in DLPFC (n=1377). The sample concentration is reported on Y axis.

**Suppl. Figure 6:** Co-immunostaining of antibodies recognizing exon 7 (red) with ALDH1L1 (green) in human *post mortem* tissue of AD and non-AD subjects. The number of astrocytes expressing the exon 7 and the total number of astrocytes have been quantified (n=5).

## Supplementary Tables

**Suppl. Table 1:** Sample demographic table for RNA-seq and SRM proteomics data.

**Suppl. Table 2:** List of 7 peptides, encoded by exons 7, 8, 10, 12 and 13 of BIN1 and selected based on peptides detectability prediction tool CONSeQuence.

**Suppl. Table 3:** List of six BIN1 peptides detected in purified human microglia, encoded by exons 15-16, 9, 5, 8, 11 and 15, all expressed in all isoforms of BIN1.

**Suppl. Table 4:** a-b: Evaluation of the effect of genetic variant on BIN1 expression at the mRNA level in human DLPFC.

**Suppl. Table 5:** Association of BIN1-7 peptides with AD dementia, cognition, and AD pathology including amyloid pathology and tangles. (fdr<0.05)

**Suppl. Table 6:** Association of BIN1-7 peptides with neurovascular measures, CAA, hippocampal sclerosis, Lewy Bodies, and the burden of TDP43 pathology. No significant association has been detected between BIN1 peptides and the pathologies listed above. (fdr<0.05)

**Suppl. Table 7:** Association of BIN1 peptides with AD pathology (amyloid pathology and tangles) has been re-analyzed using conditional analyses, by adjusting for the burden of tangles or for amyloid. (*p*<0.05)

**Suppl. Table 8:** Association of BIN1 peptides with cognitive decline by adjusting for tangles (*p*<0.05) using conditional analyses.

**Suppl. Table 9:** Analysis of the association between known AD variants and BIN1 peptides after correcting for the testing of multiple hypotheses (*p*<0.05).

**Suppl. Table 10:** Study of the association between the tau-related epigenomic alterations and BIN1 peptides (fdr<0.05)

