## supplementary figures for "BIN1 protein isoforms are differentially expressed in astrocytes, neurons, and microglia: neuronal and astrocyte BIN1 implicated in Tau pathology"

Suppl. Fig. 1

a.

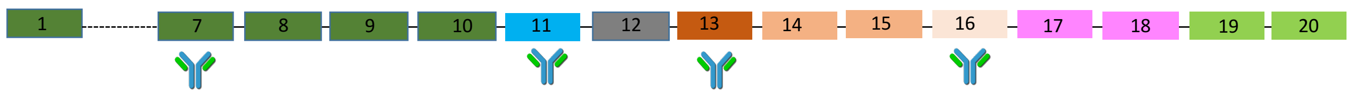

b.

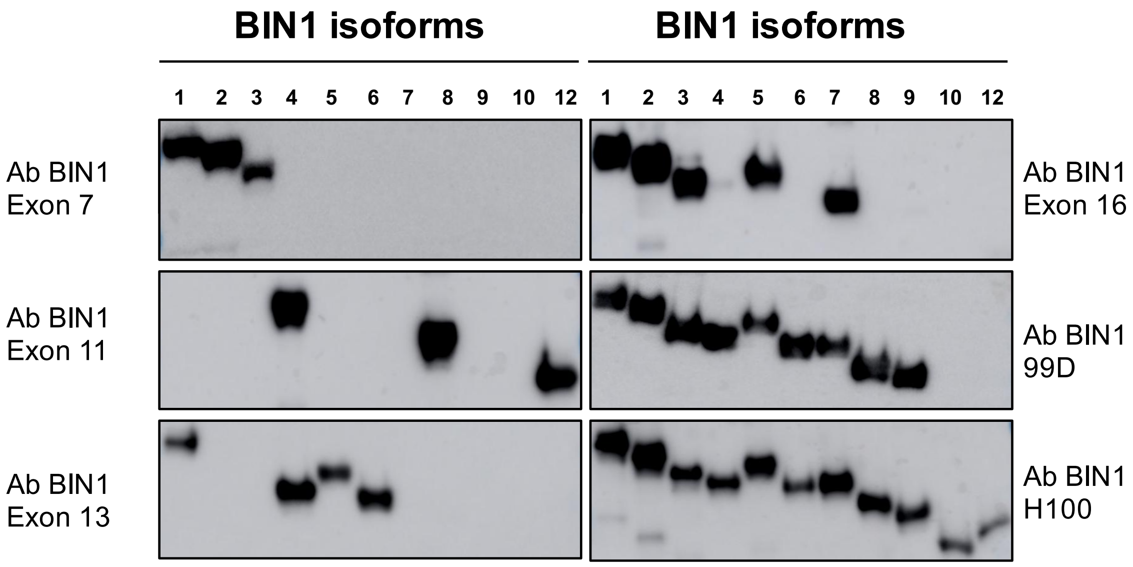

Suppl. Fig. 2

Ab BIN1 Exon 7 (1.0 µg/mL), IHC against 11 isotypes

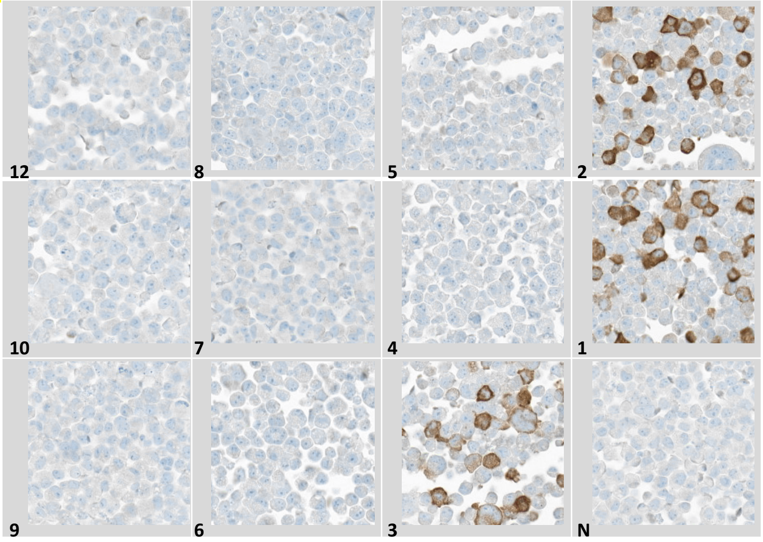

Ab BIN1 Exon 11 (0.25 µg/mL), IHCagainst 11 isotypes

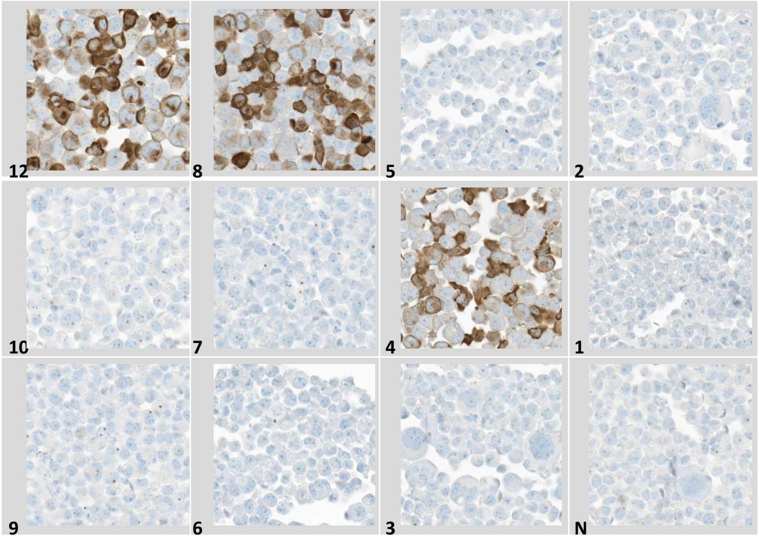

Ab BIN1 Exon 13 (1.0 µg/mL), IHC against 11 isotypes

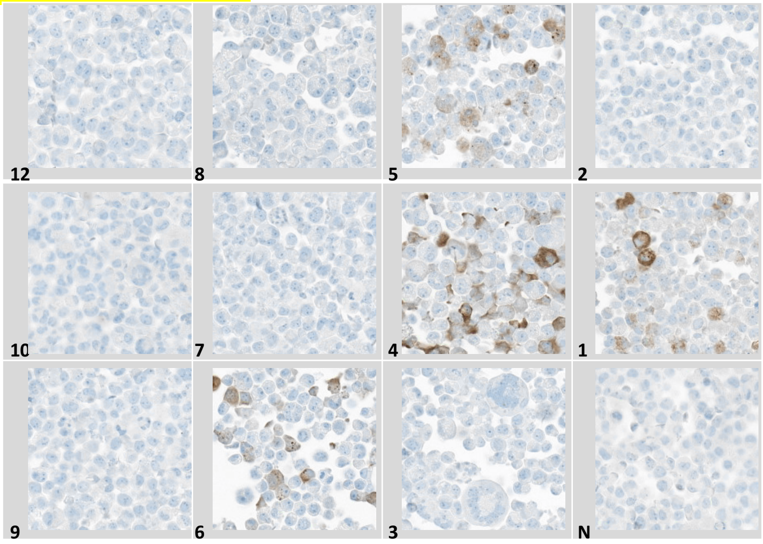

Ab BIN1 Exon 16 (0.25 µg/mL), IHCagainst 11 isotypes

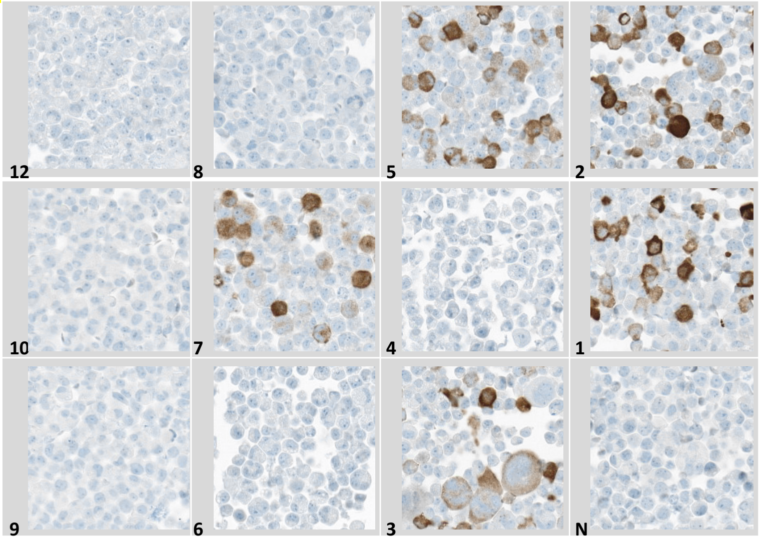

### Suppl. Fig. 3

a.

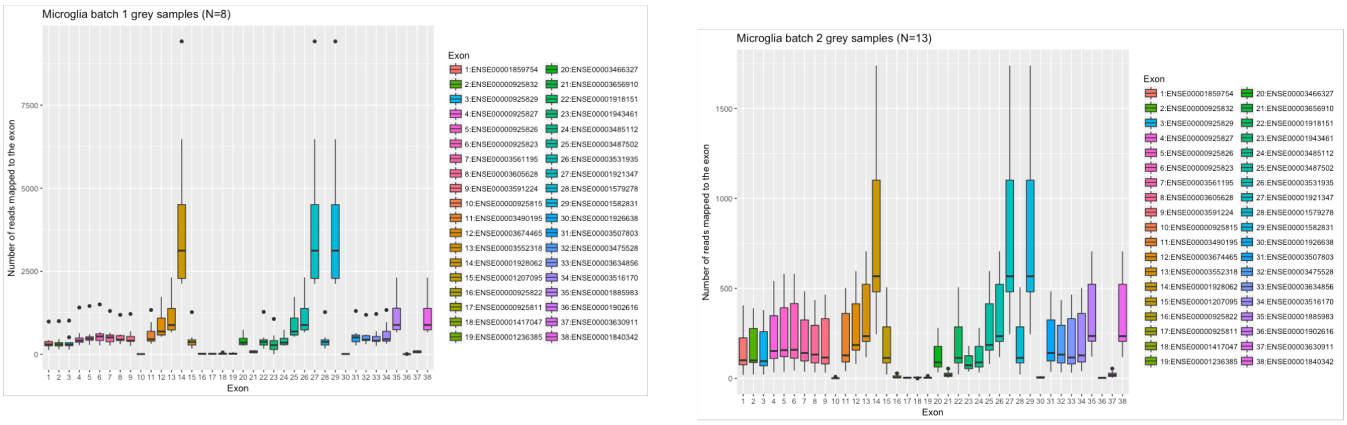

b.

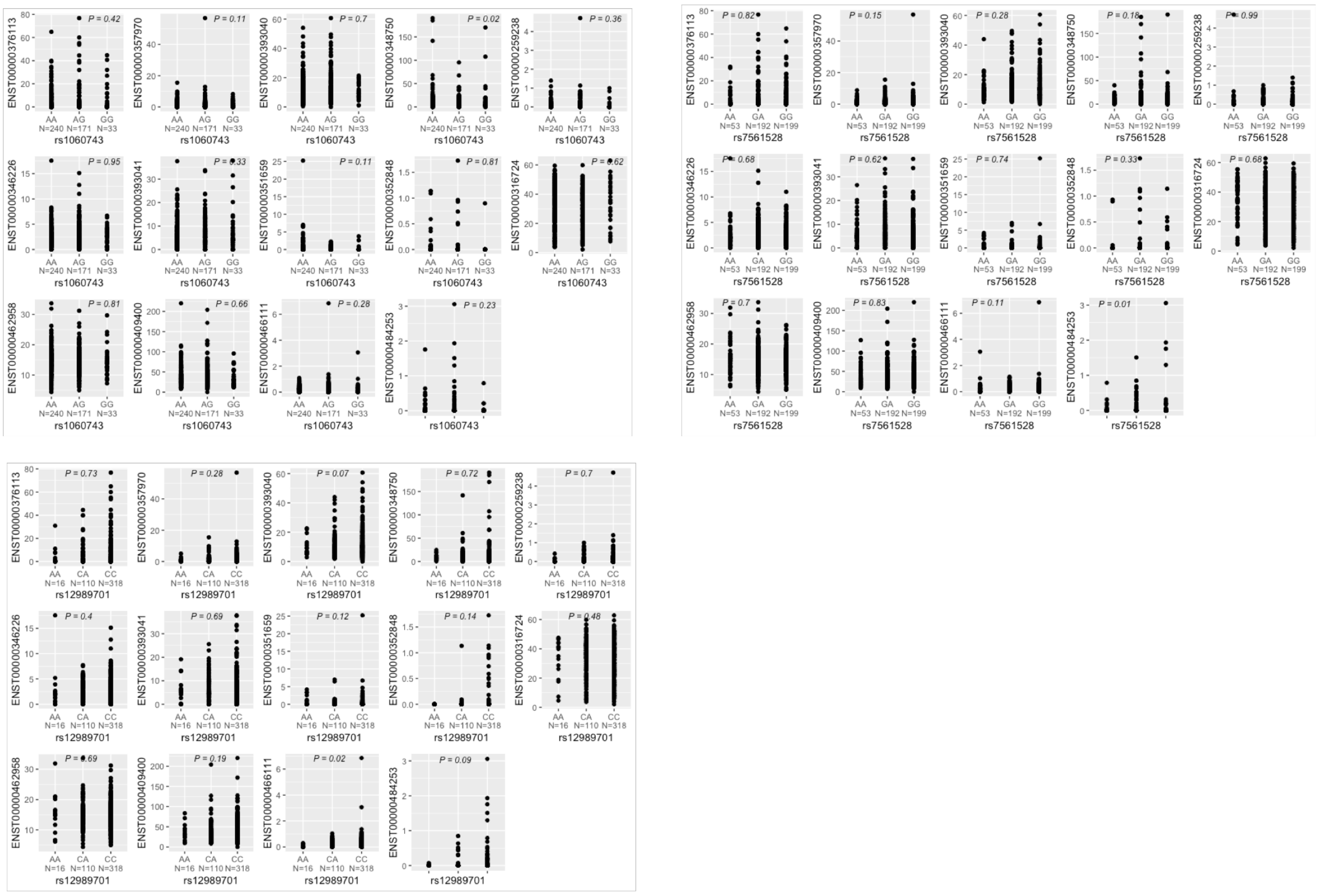

c.

| gene | snp | Value | Std.Error | DF | t.value | p.value |
|---|---|---|---|---|---|---|
| ENST00000316724 | rs1060743 | 0.02026785 | 0.03150847 | 20 | 0.64325087 | 0.52736856 |
| ENST00000346226 | rs1060743 | -0.0023177 | 0.02755779 | 20 | -0.0841025 | 0.93381107 |
| ENST00000348750 | rs1060743 | -0.5582641 | 0.50427024 | 20 | -1.1070733 | 0.28140022 |
| ENST00000351659 | rs1060743 | -0.1928578 | 0.08994637 | 20 | -2.1441425 | 0.04449238 |
| ENST00000357970 | rs1060743 | 0.00305947 | 0.01541643 | 20 | 0.19845509 | 0.84469244 |
| ENST00000376113 | rs1060743 | 0.15906423 | 0.5005467 | 20 | 0.317781 | 0.75394511 |
| ENST00000393040 | rs1060743 | -0.3192034 | 0.43230766 | 20 | -0.738371 | 0.46886564 |
| ENST00000393041 | rs1060743 | 0.12888875 | 0.08050971 | 20 | 1.60090941 | 0.12507485 |
| ENST00000409400 | rs1060743 | -0.4690434 | 0.29317641 | 20 | -1.5998676 | 0.12530648 |
| ENST00000462958 | rs1060743 | -0.2458489 | 0.19346907 | 20 | -1.2707402 | 0.21840395 |
| ENST00000466111 | rs1060743 | 0.0951159 | 0.13594505 | 20 | 0.69966433 | 0.49219538 |
| ENST00000484253 | rs1060743 | 0.1355292 | 0.15125768 | 20 | 0.8960153 | 0.38090291 |
| ENST00000316724 | rs7561528 | 0.05128569 | 0.03099093 | 20 | 1.65486132 | 0.11356155 |
| ENST00000346226 | rs7561528 | -0.00272 | 0.02887104 | 20 | -0.0942128 | 0.9258776 |
| ENST00000348750 | rs7561528 | -0.7674993 | 0.55265128 | 20 | -1.3887588 | 0.18017906 |
| ENST00000351659 | rs7561528 | -0.1405315 | 0.09905208 | 20 | -1.4187641 | 0.17136422 |
| ENST00000357970 | rs7561528 | 0.02023521 | 0.01578883 | 20 | 1.28161554 | 0.21463721 |
| ENST00000376113 | rs7561528 | 0.22671896 | 0.53766682 | 20 | 0.42167184 | 0.67776223 |
| ENST00000393040 | rs7561528 | 0.03745194 | 0.48969798 | 20 | 0.07647966 | 0.9397974 |
| ENST00000393041 | rs7561528 | -0.0291213 | 0.08747893 | 20 | -0.3328954 | 0.74267754 |
| ENST00000409400 | rs7561528 | -0.2957696 | 0.34732181 | 20 | -0.8515722 | 0.40453312 |
| ENST00000462958 | rs7561528 | -0.2182246 | 0.21825725 | 20 | -0.9998503 | 0.32932723 |
| ENST00000466111 | rs7561528 | 0.0055047 | 0.15375624 | 20 | 0.03580149 | 0.97179557 |
| ENST00000484253 | rs7561528 | 0.0271221 | 0.16318829 | 20 | 0.16620124 | 0.86966729 |
| ENST00000316724 | rs12989701 | 0.02036485 | 0.03795251 | 20 | 0.53658761 | 0.59747136 |
| ENST00000346226 | rs12989701 | -0.0314037 | 0.032609 | 20 | -0.9630372 | 0.34702919 |
| ENST00000348750 | rs12989701 | -0.4731787 | 0.64579757 | 20 | -0.7327043 | 0.47223947 |
| ENST00000351659 | rs12989701 | -0.1105369 | 0.11558904 | 20 | -0.9562927 | 0.35034183 |
| ENST00000357970 | rs12989701 | 0.00491115 | 0.01858831 | 20 | 0.26420648 | 0.79432423 |
| ENST00000376113 | rs12989701 | -0.0666061 | 0.61727135 | 20 | -0.1079041 | 0.91514697 |
| ENST00000393040 | rs12989701 | 0.24664104 | 0.54981523 | 20 | 0.44858896 | 0.65855059 |
| ENST00000393041 | rs12989701 | -0.129444 | 0.09757858 | 20 | -1.3265614 | 0.19960288 |
| ENST00000409400 | rs12989701 | -0.2631607 | 0.39401292 | 20 | -0.6678987 | 0.51183276 |
| ENST00000462958 | rs12989701 | -0.1844715 | 0.25021369 | 20 | -0.7372556 | 0.46952855 |
| ENST00000466111 | rs12989701 | -0.2038351 | 0.16371311 | 20 | -1.2450753 | 0.22749564 |
| ENST00000484253 | rs12989701 | -0.1646926 | 0.18069305 | 20 | -0.9114498 | 0.37291395 |

Suppl. Fig. 4

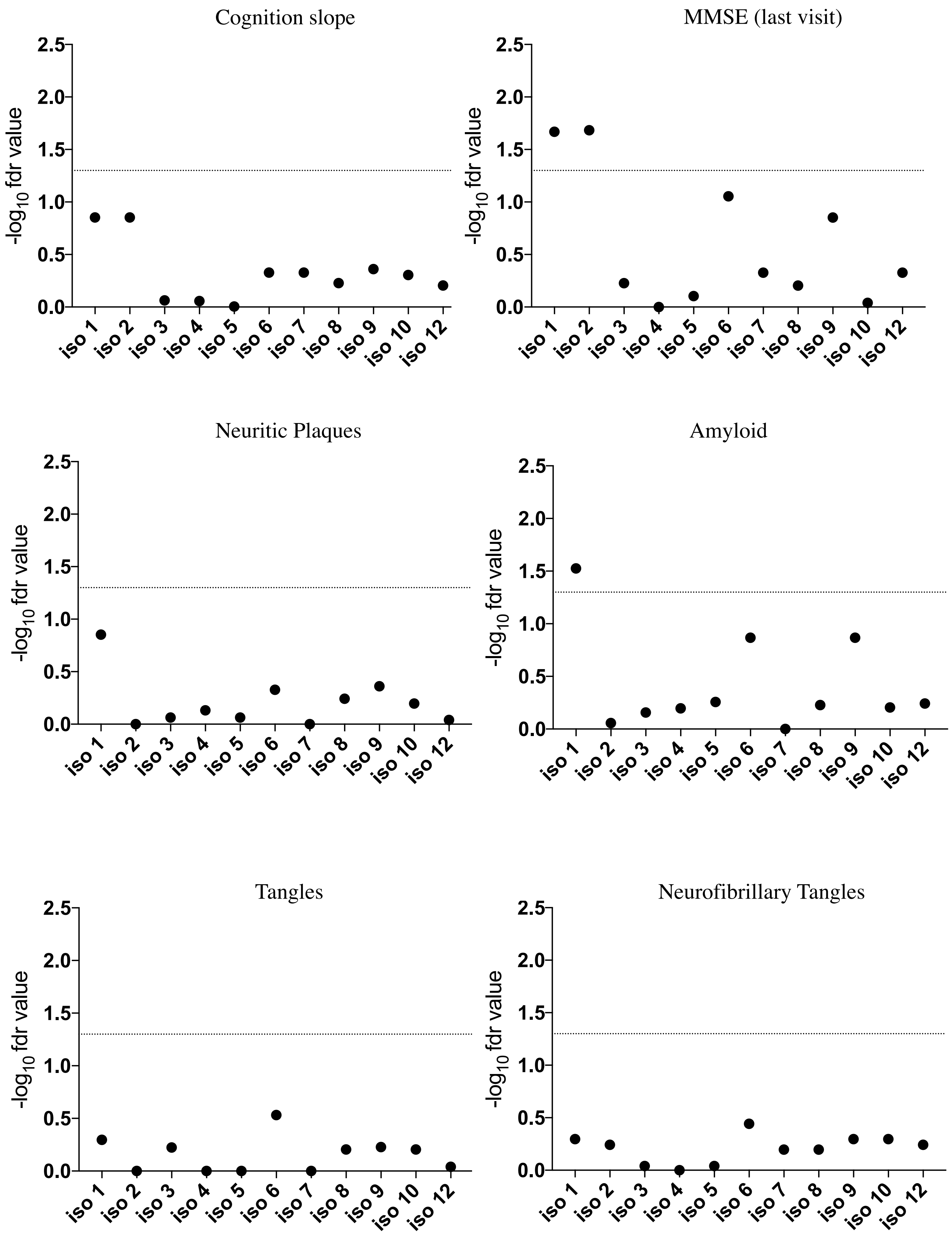
Cognition slope
MMSE (last visit)
Neuritic Plaques
Amyloid
Tangles
Neurofibrillary Tangles
$-\log_{10}$ fdr value
0.0
0.5
1.0
1.5
2.0
2.5
iso 1
iso 2
iso 3
iso 4
iso 5
iso 6
iso 7
iso 8
iso 9
iso 10
iso 12

Suppl. Fig. 5

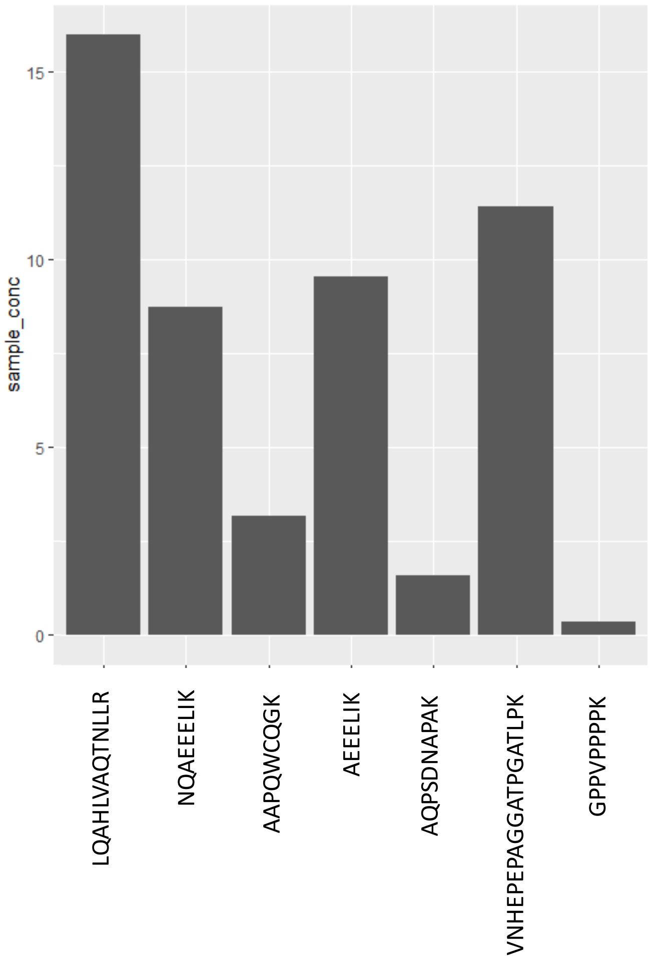
sample_conc
15
10
5
0
LQAHLVAQTNLLR
NQAEEELIK
AAPQWCQGK
AEEELIK
AQPSDNAPAK
VNHEPEPAGGATPGATLPK
GPPVPPPPK

Suppl. Fig. 6

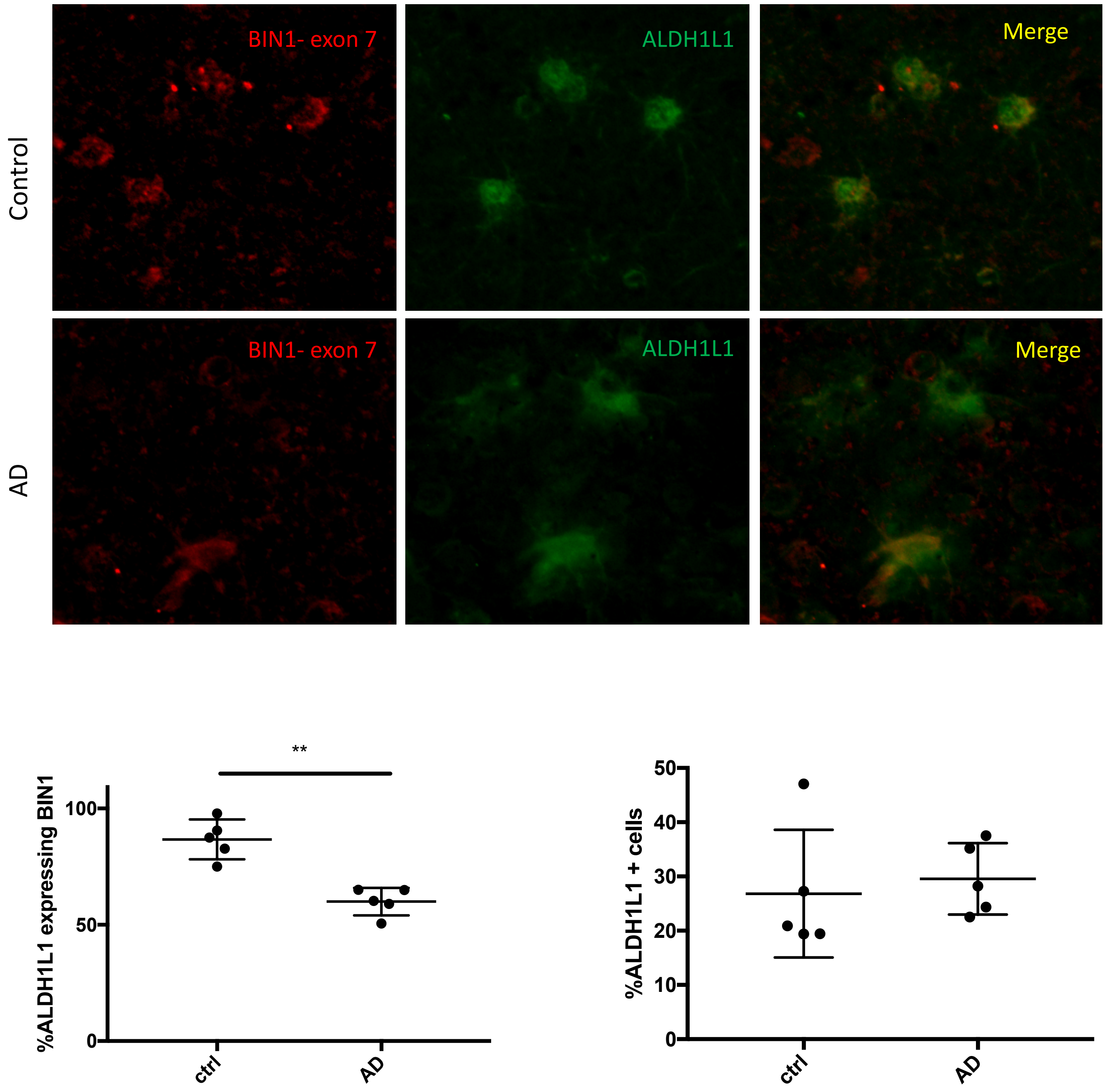
Control
AD
BIN1- exon 7
ALDH1L1
Merge
BIN1- exon 7
ALDH1L1
Merge
**
%ALDH1L1 expressing BIN1
100
50
0
ctrl
AD
%ALDH1L1 + cells
50
40
30
20
10
0
ctrl
AD

### Suppl. Table 1

| | cortical RNA-seq (n=508) |
|---|---|
| Male | 38% |
| Female | 62% |
| Age_death | 88.4 +- 6.6 |

| | SRM proteomics (n=1377) |
|---|---|
| Male | 31.70% |
| Female | 68.300% |
| Age_death | 89.4 +- 6.5 |

### Suppl. Table2

| **Peptide sequence** | **Consenquence Rank Score** | **exon** |
|---|---|---|
| LQAHLVAQTNLLR | 0.544 | 7 |
| NQAEEELIK | 0.267 | 7 |
| AAPQWCQGK | 0.255 | 7 |
| AEEELIK | 0.215 | 8 |
| AQPSDNAPAK | 0.236 | 10 |
| VNHEPEPAGGATPGATLPK | 0.899 | 12 |
| GPPVPPPPK | 0.085 | 13 |

### Suppl. Table 3

| **Peptide sequence** | **exon** | **isoform** |
|---|---|---|
| AGDVVLVIPFQNPEEQDEGWLMGV | 15-16 | all isoform |
| LNQNLNDVLVGLEK | 9 | all isoform |
| LVDQALLTMDTYLGQFPDIK | 5 | all isoform |
| VGFYVNTFQSIAGLEENFHK | 8 | all isoform |
| VNHEPEPAGGATPGATLPK | 11 | all isoform |
| VQAQHDYTATDTDELQLK | 15 | all isoform |

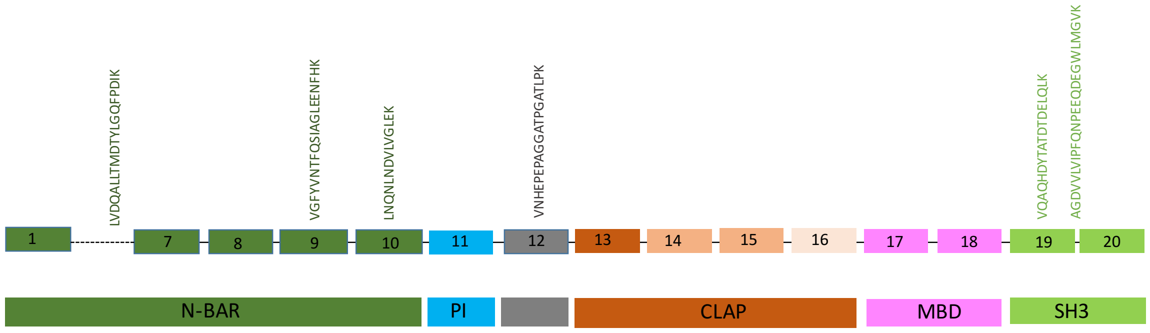

### Suppl. Table 4

a.

| BIN1 isoforms | ENST_gencode19 | number of SNPs < 5.57E-5 | number of SNPs < 0.05 |
|---|---|---|---|
| 1 | ENST00000316724 | 0 | 73 |
| 2 | ENST00000357970 | 0 | 6 |
| 3 | ENST00000351659 | 0 | 194 |
| 4 | ENST00000259238 | 0 | 3 |
| 5 | ENST00000346226 | 0 | 0 |
| 6 | ENST00000393040 | 0 | 65 |
| 7 | ENST00000393041 | 0 | 127 |
| 8 | ENST00000352848 | N/A | N/A |
| 9 | ENST00000409400 | 0 | 60 |
| 10 | ENST00000348750 | 0 | 82 |
| 12 | ENST00000376113 | 0 | 57 |

b.

| Id | Gencode_id | chr | position | Effective allele | a2 | mac | Freq | Effect | Stdev | Z | *p* |
|---|---|---|---|---|---|---|---|---|---|---|---|
| rs1060743 | ENST00000348750.4 | 10,2 | 127826533 | A | G | 262.96 | 0.73 | -0.16 | 0.07 | -2.25 | 0.02 |

#### Suppl. Table 5

| Peptides | *fdr* _AD dementia | *fdr* _Slope of Cognitive Decline | *fdr* _Residual Cog | *fdr* _Pathologic Diag. | *fdr* _Amyloid Patho | *fdr* _tangles |
|---|---|---|---|---|---|---|
| **LQAHLVAQTNLLR** | 0.026007692 | 0.0002115 | 0.000780938 | 0.0002954 | 1.91E-05 | 1.89E-07 |
| **NQAEEELIK** | 0.001569167 | 6.98E-05 | 1.16E-05 | 6.84E-05 | 6.26E-08 | 2.34E-09 |
| **AAPQWCQGK** | 0.112397727 | 0.027375 | 0.026007692 | 0.00358575 | 0.052181818 | 0.00329921 |
| **AEEELIK** | 0.553875 | 0.523561644 | 0.290625 | 0.253584906 | 0.013923913 | 0.28254545 |
| **AQPSDNAPAK** | 0.770823529 | 0.544090909 | 0.318559322 | 0.318559322 | 0.01435 | 0.11239773 |
| **VNHEPEPAGGATPGATLPK** | 0.907644231 | 0.811774194 | 0.523561644 | 0.527837838 | 0.1579375 | 0.54759494 |
| **GPPVPPPPK** | 0.122266667 | 0.028965517 | 0.026911111 | 0.434538462 | 0.907644231 | 0.11239773 |

#### Suppl. Table 6

| Peptides | *fdr* _CAA | *fdr* _ci_num2_gct | *fdr* _ci_num2_mct | *fdr* _Lew Body | *fdr* _Hippocampal Sclerosis | *fdr* _TDP43 pathology |
|---|---|---|---|---|---|---|
| **LQAHLVAQTNLLR** | 0.033396774 | 0.929 | 0.845585106 | 0.875357143 | 0.055897059 | 0.156382979 |
| **NQAEEELIK** | 0.033396774 | 0.774375 | 0.888787879 | 0.7425 | 0.16632 | 0.16632 |
| **AAPQWCQGK** | 0.070291667 | 0.875357143 | 0.771627907 | 0.875357143 | 0.38675 | 0.501338028 |
| **AEEELIK** | 0.544090909 | 0.907644231 | 0.807333333 | 0.697048193 | 0.656890244 | 0.501338028 |
| **AQPSDNAPAK** | 0.849947368 | 0.076263158 | 0.547594937 | 0.907644231 | 0.58462963 | 0.791629213 |
| **VNHEPEPAGGATPGATLPK** | 0.470149254 | 0.811774194 | 0.774375 | 0.397622951 | 0.811774194 | 0.486397059 |
| **GPPVPPPPK** | 0.09555 | 0.907644231 | 0.544090909 | 0.172941176 | 0.451818182 | 0.434538462 |

#### Suppl. Table 7

| Peptides | *fdr* _AD dementia | *fdr* _Slope of Cognitive Decline | *fdr* _Residual Cog | *fdr* _Pathologic Diag. | *fdr* _Amyloid Patho | *fdr* _tangles |
|---|---|---|---|---|---|---|
| **LQAHLVAQTNLLR** | 0.833269231 | 0.159 | 0.2905 | 0.036 | 0.0105 | 0.02044875 |
| **NQAEEELIK** | 0.853915663 | 0.7940625 | 0.792105263 | 0.036 | 0.02044875 | 0.389454545 |
| **AAPQWCQGK** | 0.512195122 | 0.448636364 | 0.448636364 | 0.118363636 | 0.344842105 | 0.285352941 |
| **AEEELIK** | 0.920684211 | 0.853915663 | 0.823636364 | 0.448636364 | 0.792105263 | 0.9576 |
| **AQPSDNAPAK** | 0.448636364 | 0.777 | 0.719150943 | 0.862670455 | 0.823636364 | 0.495833333 |
| **VNHEPEPAGGATPGATLPK** | 0.823636364 | 0.823636364 | 0.823636364 | 0.63 | 0.890217391 | 0.853915663 |
| **GPPVPPPPK** | 0.823636364 | 0.525 | 0.495833333 | 0.512195122 | 0.7940625 | 0.814545455 |

#### Suppl. Table 8

| Peptides | *p* _Amyloid_adj | *p*_ Tau_adj |
|---|---|---|
| AAPQWCQGK | 0.32 | 2.58E-05 |
| LQAHLVAQTNLLR | 0.04 | 2.11E-06 |
| NQAEEELIK | 0.000762 | 3.40E-05 |
| AEEELIK | 0.0323 | 0.72 |
| AQPSDNAPAK | 0.0447 | 0.469 |
| VNHEPEPAGGATPGATLPK | 0.103 | 0.849 |
| GPPVPPPPK | 0.357 | 0.0134 |

#### Suppl. Table 9

| Peptide | Beta_peptide_cogdec | R2_peptide_cogdec | *p* _peptide_cogdec |
|---|---|---|---|
| **LQAHLVAQTNLLR** | 0.0564 | 0.0148 | 4.63E-05 |
| **NQAEEELIK** | 0.0775 | 0.0179 | 7.34E-06 |
| **AAPQWCQGK** | 0.0287 | 0.00553 | 0.013 |
| **AEEELIK** | -0.00473 | 0.00201 | 0.134 |
| **AQPSDNAPAK** | -0.00497 | 0.00117 | 0.254 |
| **VNHEPEPAGGATPGATLPK** | -0.00547 | 0.000736 | 0.365 |
| **GPPVPPPPK** | 0.0262 | 0.00315 | 0.0605 |

| Beta_tau_cogdec | R2_tangles_cogdec | *p* _tangles_cogdec |
|---|---|---|
| -0.0327 | 0.2 | 7.59E-67 |

| Peptide | Beta_peptide_cogdec_jt | R2_peptide_cogdec_jt | p_peptide_cogdec_jt | Beta_tangles_cogdec_jt | R2_tangles_cogdec_jt | p_tangles_cogdec_jt |
|---|---|---|---|---|---|---|
| **LQAHLVAQTNLLR** | 0.029 | 0.00488 | 0.0202 | -0.0332 | 0.197 | 1.59E-54 |
| **NQAEEELIK** | 0.0377 | 0.00531 | 0.0153 | -0.0331 | 0.196 | 3.22E-54 |
| **AAPQWCQGK** | 0.0138 | 0.00163 | 0.18 | -0.0333 | 0.199 | 5.36E-55 |
| **AEEELIK** | -0.0047 | 0.00256 | 0.0927 | -0.0339 | 0.207 | 1.65E-57 |
| **AQPSDNAPAK** | -0.00404 | 0.000999 | 0.295 | -0.0337 | 0.204 | 2.98E-56 |
| **VNHEPEPAGGATPGATLPK** | -0.0063 | 0.00126 | 0.238 | -0.0339 | 0.207 | 1.99E-57 |
| **GPPVPPPPK** | 0.00697 | 0.000285 | 0.575 | -0.0338 | 0.204 | 1.07E-56 |

### Suppl. Table 10

| SNP | Peptides | beta | se | tstat | pval | n | df |
|---|---|---|---|---|---|---|---|
| rs10498633 | LQAHLVAQTNLLR | -0.0022 | 0.0114 | -0.194 | 0.846 | 1018 | 5;1013 |
| rs10498633 | NQAEEELIK | -0.00343 | 0.00893 | -0.384 | 0.701 | 1018 | 5;1013 |
| rs10498633 | AAPQWCQGK | -0.0111 | 0.0127 | -0.876 | 0.381 | 1015 | 5;1010 |
| rs10498633 | AEEELIK | -0.0257 | 0.0505 | -0.508 | 0.611 | 1018 | 5;1013 |
| rs10498633 | AQPSDNAPAK | 0.00223 | 0.0368 | 0.0607 | 0.952 | 1012 | 5;1007 |
| rs10498633 | VNHEPEPAGGATPGATLPK | -0.0137 | 0.0265 | -0.517 | 0.605 | 1017 | 5;1012 |
| rs10498633 | GPPVPPPPK | -0.0055 | 0.0113 | -0.485 | 0.628 | 1018 | 5;1013 |
| rs10792832 | LQAHLVAQTNLLR | 0.00533 | 0.01 | 0.531 | 0.596 | 1018 | 5;1013 |
| rs10792832 | NQAEEELIK | 0.0147 | 0.00788 | 1.86 | 0.0631 | 1018 | 5;1013 |
| rs10792832 | AAPQWCQGK | -0.00608 | 0.0112 | -0.542 | 0.588 | 1015 | 5;1010 |
| rs10792832 | AEEELIK | -0.0803 | 0.0446 | -1.8 | 0.0722 | 1018 | 5;1013 |
| rs10792832 | AQPSDNAPAK | -0.0128 | 0.0325 | -0.395 | 0.693 | 1012 | 5;1007 |
| rs10792832 | VNHEPEPAGGATPGATLPK | -0.0286 | 0.0234 | -1.22 | 0.222 | 1017 | 5;1012 |
| rs10792832 | GPPVPPPPK | -0.0152 | 0.01 | -1.52 | 0.129 | 1018 | 5;1013 |
| rs10838725 | LQAHLVAQTNLLR | -0.00383 | 0.0105 | -0.364 | 0.716 | 1018 | 5;1013 |
| rs10838725 | NQAEEELIK | -0.0026 | 0.00827 | -0.314 | 0.753 | 1018 | 5;1013 |
| rs10838725 | AAPQWCQGK | -0.0124 | 0.0117 | -1.05 | 0.293 | 1015 | 5;1010 |
| rs10838725 | AEEELIK | 0.0197 | 0.0468 | 0.42 | 0.675 | 1018 | 5;1013 |
| rs10838725 | AQPSDNAPAK | -0.0263 | 0.034 | -0.773 | 0.44 | 1012 | 5;1007 |
| rs10838725 | VNHEPEPAGGATPGATLPK | 0.0304 | 0.0245 | 1.24 | 0.216 | 1017 | 5;1012 |
| rs10838725 | GPPVPPPPK | 0.00472 | 0.0105 | 0.449 | 0.653 | 1018 | 5;1013 |
| rs10948363 | LQAHLVAQTNLLR | -0.0192 | 0.0109 | -1.76 | 0.0792 | 1018 | 5;1013 |
| rs10948363 | NQAEEELIK | -0.0196 | 0.0086 | -2.27 | 0.0231 | 1018 | 5;1013 |
| rs10948363 | AAPQWCQGK | -0.0136 | 0.0122 | -1.11 | 0.268 | 1015 | 5;1010 |
| rs10948363 | AEEELIK | 0.0223 | 0.0488 | 0.456 | 0.649 | 1018 | 5;1013 |
| rs10948363 | AQPSDNAPAK | 0.00882 | 0.0356 | 0.248 | 0.804 | 1012 | 5;1007 |
| rs10948363 | VNHEPEPAGGATPGATLPK | 0.0139 | 0.0256 | 0.543 | 0.587 | 1017 | 5;1012 |
| rs10948363 | GPPVPPPPK | -0.00565 | 0.0109 | -0.517 | 0.606 | 1018 | 5;1013 |
| rs11218343 | LQAHLVAQTNLLR | -0.00627 | 0.0267 | -0.234 | 0.815 | 1018 | 5;1013 |
| rs11218343 | NQAEEELIK | -0.00969 | 0.021 | -0.461 | 0.645 | 1018 | 5;1013 |
| rs11218343 | AAPQWCQGK | -0.0192 | 0.0298 | -0.643 | 0.52 | 1015 | 5;1010 |
| rs11218343 | AEEELIK | 0.0126 | 0.119 | 0.106 | 0.916 | 1018 | 5;1013 |
| rs11218343 | AQPSDNAPAK | -0.0237 | 0.087 | -0.273 | 0.785 | 1012 | 5;1007 |
| rs11218343 | VNHEPEPAGGATPGATLPK | 0.0106 | 0.0624 | 0.17 | 0.865 | 1017 | 5;1012 |
| rs11218343 | GPPVPPPPK | -0.000202 | 0.0267 | -0.00758 | 0.994 | 1018 | 5;1013 |
| rs1171812 | LQAHLVAQTNLLR | 0.00831 | 0.0095 | 0.875 | 0.382 | 1018 | 5;1013 |
| rs1171812 | NQAEEELIK | 0.0132 | 0.00747 | 1.77 | 0.0776 | 1018 | 5;1013 |
| rs1171812 | AAPQWCQGK | 0.0216 | 0.0106 | 2.04 | 0.0419 | 1015 | 5;1010 |
| rs1171812 | AEEELIK | -0.0295 | 0.0423 | -0.697 | 0.486 | 1018 | 5;1013 |
| rs1171812 | AQPSDNAPAK | -0.00562 | 0.0308 | -0.182 | 0.855 | 1012 | 5;1007 |
| rs1171812 | VNHEPEPAGGATPGATLPK | -0.0131 | 0.0222 | -0.589 | 0.556 | 1017 | 5;1012 |
| rs1171812 | GPPVPPPPK | 0.00985 | 0.00949 | 1.04 | 0.3 | 1018 | 5;1013 |
| rs11771145 | LQAHLVAQTNLLR | 0.00532 | 0.0114 | 0.465 | 0.642 | 1018 | 5;1013 |
| rs11771145 | NQAEEELIK | 0.00694 | 0.009 | 0.771 | 0.441 | 1018 | 5;1013 |
| rs11771145 | AAPQWCQGK | 0.00109 | 0.0128 | 0.0849 | 0.932 | 1015 | 5;1010 |
| rs11771145 | AEEELIK | -0.0856 | 0.0509 | -1.68 | 0.093 | 1018 | 5;1013 |
| rs11771145 | AQPSDNAPAK | -0.037 | 0.0371 | -0.996 | 0.319 | 1012 | 5;1007 |
| rs11771145 | VNHEPEPAGGATPGATLPK | -0.0396 | 0.0267 | -1.49 | 0.138 | 1017 | 5;1012 |
| rs11771145 | GPPVPPPPK | 0.00982 | 0.0114 | 0.86 | 0.39 | 1018 | 5;1013 |
| rs12444183 | LQAHLVAQTNLLR | 0.00205 | 0.01 | 0.205 | 0.838 | 1018 | 5;1013 |
| rs12444183 | NQAEEELIK | -0.0043 | 0.00789 | -0.545 | 0.586 | 1018 | 5;1013 |
| rs12444183 | AAPQWCQGK | -0.0105 | 0.0112 | -0.935 | 0.35 | 1015 | 5;1010 |
| rs12444183 | AEEELIK | 0.0392 | 0.0447 | 0.878 | 0.38 | 1018 | 5;1013 |
| rs12444183 | AQPSDNAPAK | -0.0285 | 0.0325 | -0.877 | 0.381 | 1012 | 5;1007 |
| rs12444183 | VNHEPEPAGGATPGATLPK | 0.0213 | 0.0234 | 0.909 | 0.364 | 1017 | 5;1012 |
| rs12444183 | GPPVPPPPK | -0.000817 | 0.01 | -0.0815 | 0.935 | 1018 | 5;1013 |
| rs1476679 | LQAHLVAQTNLLR | -0.0172 | 0.0105 | -1.64 | 0.1 | 1018 | 5;1013 |
| rs1476679 | NQAEEELIK | 0.00479 | 0.00824 | 0.581 | 0.561 | 1018 | 5;1013 |
| rs1476679 | AAPQWCQGK | -0.00233 | 0.0117 | -0.199 | 0.842 | 1015 | 5;1010 |
| rs1476679 | AEEELIK | -0.0192 | 0.0466 | -0.411 | 0.681 | 1018 | 5;1013 |
| rs1476679 | AQPSDNAPAK | -0.0504 | 0.0339 | -1.49 | 0.138 | 1012 | 5;1007 |
| rs1476679 | VNHEPEPAGGATPGATLPK | -0.017 | 0.0245 | -0.695 | 0.487 | 1017 | 5;1012 |
| rs1476679 | GPPVPPPPK | 0.000948 | 0.0105 | 0.0907 | 0.928 | 1018 | 5;1013 |
| rs17125944 | LQAHLVAQTNLLR | -0.016 | 0.0169 | -0.95 | 0.342 | 1018 | 5;1013 |
| rs17125944 | NQAEEELIK | -0.00259 | 0.0133 | -0.195 | 0.845 | 1018 | 5;1013 |
| rs17125944 | AAPQWCQGK | -0.00728 | 0.0188 | -0.387 | 0.699 | 1015 | 5;1010 |
| rs17125944 | AEEELIK | 0.0515 | 0.0751 | 0.686 | 0.493 | 1018 | 5;1013 |
| rs17125944 | AQPSDNAPAK | 0.0141 | 0.0547 | 0.258 | 0.797 | 1012 | 5;1007 |
| rs17125944 | VNHEPEPAGGATPGATLPK | 0.00547 | 0.0393 | 0.139 | 0.89 | 1017 | 5;1012 |
| rs17125944 | GPPVPPPPK | 0.00316 | 0.0168 | 0.188 | 0.851 | 1018 | 5;1013 |
| rs190982 | LQAHLVAQTNLLR | 0.00709 | 0.0106 | 0.668 | 0.504 | 1018 | 5;1013 |
| rs190982 | NQAEEELIK | -0.00208 | 0.00835 | -0.249 | 0.804 | 1018 | 5;1013 |
| rs190982 | AAPQWCQGK | 0.00266 | 0.0119 | 0.224 | 0.823 | 1015 | 5;1010 |
| rs190982 | AEEELIK | -0.0298 | 0.0473 | -0.63 | 0.529 | 1018 | 5;1013 |
| rs190982 | AQPSDNAPAK | -0.0178 | 0.0344 | -0.518 | 0.605 | 1012 | 5;1007 |
| rs190982 | VNHEPEPAGGATPGATLPK | -0.0136 | 0.0248 | -0.551 | 0.582 | 1017 | 5;1012 |
| rs190982 | GPPVPPPPK | 0.00721 | 0.0106 | 0.68 | 0.497 | 1018 | 5;1013 |
| rs2718058 | LQAHLVAQTNLLR | -0.0198 | 0.0105 | -1.89 | 0.0586 | 1018 | 5;1013 |
| rs2718058 | NQAEEELIK | -0.00316 | 0.00826 | -0.382 | 0.702 | 1018 | 5;1013 |
| rs2718058 | AAPQWCQGK | -0.000266 | 0.0118 | -0.0227 | 0.982 | 1015 | 5;1010 |
| rs2718058 | AEEELIK | -0.0293 | 0.0468 | -0.626 | 0.531 | 1018 | 5;1013 |
| rs2718058 | AQPSDNAPAK | -0.0104 | 0.0341 | -0.304 | 0.761 | 1012 | 5;1007 |
| rs2718058 | VNHEPEPAGGATPGATLPK | -0.0157 | 0.0245 | -0.643 | 0.521 | 1017 | 5;1012 |
| rs2718058 | GPPVPPPPK | -0.00641 | 0.0105 | -0.612 | 0.541 | 1018 | 5;1013 |
| rs28834970 | LQAHLVAQTNLLR | -0.0128 | 0.0101 | -1.26 | 0.207 | 1018 | 5;1013 |
| rs28834970 | NQAEEELIK | 0.00872 | 0.00796 | 1.1 | 0.273 | 1018 | 5;1013 |
| rs28834970 | AAPQWCQGK | 0.0234 | 0.0113 | 2.08 | 0.0379 | 1015 | 5;1010 |
| rs28834970 | AEEELIK | 0.0317 | 0.0451 | 0.704 | 0.481 | 1018 | 5;1013 |
| rs28834970 | AQPSDNAPAK | 0.0325 | 0.0327 | 0.992 | 0.322 | 1012 | 5;1007 |
| rs28834970 | VNHEPEPAGGATPGATLPK | 0.0175 | 0.0236 | 0.742 | 0.458 | 1017 | 5;1012 |
| rs28834970 | GPPVPPPPK | 0.0196 | 0.0101 | 1.94 | 0.0522 | 1018 | 5;1013 |

| SNP | Peptides | beta | se | tstat | pval | n | df |
|---|---|---|---|---|---|---|---|
| rs35349669 | LQAHLVAQTNLLR | -0.00368 | 0.0107 | -0.343 | 0.732 | 1018 | 5;1013 |
| rs35349669 | NQAEEELIK | -0.00533 | 0.00844 | -0.631 | 0.528 | 1018 | 5;1013 |
| rs35349669 | AAPQWCQGK | 0.000858 | 0.012 | 0.0716 | 0.943 | 1015 | 5;1010 |
| rs35349669 | AEEELIK | 0.0186 | 0.0478 | 0.389 | 0.697 | 1018 | 5;1013 |
| rs35349669 | AQPSDNAPAK | 0.000725 | 0.0348 | 0.0208 | 0.983 | 1012 | 5;1007 |
| rs35349669 | VNHEPEPAGGATPGATLPK | 0.00949 | 0.025 | 0.379 | 0.705 | 1017 | 5;1012 |
| rs35349669 | GPPVPPPPK | 0.00798 | 0.0107 | 0.745 | 0.457 | 1018 | 5;1013 |
| rs3865444 | LQAHLVAQTNLLR | -0.00215 | 0.0105 | -0.206 | 0.837 | 1018 | 5;1013 |
| rs3865444 | NQAEEELIK | -0.00322 | 0.00823 | -0.391 | 0.696 | 1018 | 5;1013 |
| rs3865444 | AAPQWCQGK | -0.0178 | 0.0117 | -1.53 | 0.127 | 1015 | 5;1010 |
| rs3865444 | AEEELIK | -0.0246 | 0.0466 | -0.528 | 0.598 | 1018 | 5;1013 |
| rs3865444 | AQPSDNAPAK | 0.0219 | 0.0339 | 0.647 | 0.518 | 1012 | 5;1007 |
| rs3865444 | VNHEPEPAGGATPGATLPK | -0.00685 | 0.0244 | -0.281 | 0.779 | 1017 | 5;1012 |
| rs3865444 | GPPVPPPPK | -0.0191 | 0.0104 | -1.83 | 0.0676 | 1018 | 5;1013 |
| rs4147929 | LQAHLVAQTNLLR | -0.0187 | 0.0141 | -1.32 | 0.186 | 1018 | 5;1013 |
| rs4147929 | NQAEEELIK | -0.000184 | 0.0111 | -0.0166 | 0.987 | 1018 | 5;1013 |
| rs4147929 | AAPQWCQGK | -0.00561 | 0.0158 | -0.355 | 0.722 | 1015 | 5;1010 |
| rs4147929 | AEEELIK | 0.032 | 0.0629 | 0.508 | 0.612 | 1018 | 5;1013 |
| rs4147929 | AQPSDNAPAK | 0.052 | 0.0457 | 1.14 | 0.256 | 1012 | 5;1007 |
| rs4147929 | VNHEPEPAGGATPGATLPK | 0.0484 | 0.033 | 1.47 | 0.142 | 1017 | 5;1012 |
| rs4147929 | GPPVPPPPK | -0.00465 | 0.0141 | -0.329 | 0.742 | 1018 | 5;1013 |
| rs442495 | LQAHLVAQTNLLR | -0.00222 | 0.0105 | -0.212 | 0.832 | 1018 | 5;1013 |
| rs442495 | NQAEEELIK | -0.00994 | 0.00822 | -1.21 | 0.227 | 1018 | 5;1013 |
| rs442495 | AAPQWCQGK | -0.0241 | 0.0117 | -2.07 | 0.0388 | 1015 | 5;1010 |
| rs442495 | AEEELIK | 0.0153 | 0.0466 | 0.328 | 0.743 | 1018 | 5;1013 |
| rs442495 | AQPSDNAPAK | -0.0106 | 0.0339 | -0.313 | 0.754 | 1012 | 5;1007 |
| rs442495 | VNHEPEPAGGATPGATLPK | 0.00542 | 0.0244 | 0.222 | 0.824 | 1017 | 5;1012 |
| rs442495 | GPPVPPPPK | -0.0144 | 0.0104 | -1.38 | 0.169 | 1018 | 5;1013 |
| rs4575098 | LQAHLVAQTNLLR | -0.00354 | 0.0119 | -0.297 | 0.767 | 1018 | 5;1013 |
| rs4575098 | NQAEEELIK | 0.00227 | 0.00938 | 0.241 | 0.809 | 1018 | 5;1013 |
| rs4575098 | AAPQWCQGK | 0.015 | 0.0133 | 1.12 | 0.261 | 1015 | 5;1010 |
| rs4575098 | AEEELIK | -0.0148 | 0.0531 | -0.278 | 0.781 | 1018 | 5;1013 |
| rs4575098 | AQPSDNAPAK | 0.0371 | 0.0386 | 0.96 | 0.337 | 1012 | 5;1007 |
| rs4575098 | VNHEPEPAGGATPGATLPK | -0.0191 | 0.0278 | -0.687 | 0.492 | 1017 | 5;1012 |
| rs4575098 | GPPVPPPPK | -0.000825 | 0.0119 | -0.0693 | 0.945 | 1018 | 5;1013 |
| rs6448453 | LQAHLVAQTNLLR | 0.0274 | 0.0108 | 2.53 | 0.0115 | 1018 | 5;1013 |
| rs6448453 | NQAEEELIK | 0.00999 | 0.00852 | 1.17 | 0.241 | 1018 | 5;1013 |
| rs6448453 | AAPQWCQGK | 0.00987 | 0.0121 | 0.815 | 0.415 | 1015 | 5;1010 |
| rs6448453 | AEEELIK | -0.0102 | 0.0483 | -0.211 | 0.833 | 1018 | 5;1013 |
| rs6448453 | AQPSDNAPAK | 0.0111 | 0.0351 | 0.317 | 0.751 | 1012 | 5;1007 |
| rs6448453 | VNHEPEPAGGATPGATLPK | -0.00807 | 0.0253 | -0.319 | 0.75 | 1017 | 5;1012 |
| rs6448453 | GPPVPPPPK | 0.00233 | 0.0108 | 0.215 | 0.829 | 1018 | 5;1013 |
| rs6504163 | LQAHLVAQTNLLR | 0.00948 | 0.0117 | 0.811 | 0.418 | 1018 | 5;1013 |
| rs6504163 | NQAEEELIK | 0.0145 | 0.00918 | 1.58 | 0.115 | 1018 | 5;1013 |
| rs6504163 | AAPQWCQGK | 0.0136 | 0.0131 | 1.04 | 0.297 | 1015 | 5;1010 |
| rs6504163 | AEEELIK | 0.0215 | 0.0521 | 0.413 | 0.68 | 1018 | 5;1013 |
| rs6504163 | AQPSDNAPAK | -0.0503 | 0.0379 | -1.33 | 0.185 | 1012 | 5;1007 |
| rs6504163 | VNHEPEPAGGATPGATLPK | 0.00355 | 0.0273 | 0.13 | 0.897 | 1017 | 5;1012 |
| rs6504163 | GPPVPPPPK | -0.00336 | 0.0117 | -0.288 | 0.774 | 1018 | 5;1013 |
| rs6656401 | LQAHLVAQTNLLR | 0.0149 | 0.0127 | 1.17 | 0.241 | 1018 | 5;1013 |
| rs6656401 | NQAEEELIK | 0.0107 | 0.00997 | 1.07 | 0.283 | 1018 | 5;1013 |
| rs6656401 | AAPQWCQGK | 0.0218 | 0.0142 | 1.54 | 0.124 | 1015 | 5;1010 |
| rs6656401 | AEEELIK | -0.0426 | 0.0565 | -0.755 | 0.45 | 1018 | 5;1013 |
| rs6656401 | AQPSDNAPAK | -0.0377 | 0.0413 | -0.912 | 0.362 | 1012 | 5;1007 |
| rs6656401 | VNHEPEPAGGATPGATLPK | -0.0185 | 0.0296 | -0.624 | 0.533 | 1017 | 5;1012 |
| rs6656401 | GPPVPPPPK | 0.0157 | 0.0127 | 1.24 | 0.215 | 1018 | 5;1013 |
| rs6733839 | LQAHLVAQTNLLR | 0.0155 | 0.0125 | 1.24 | 0.217 | 1018 | 5;1013 |
| rs6733839 | NQAEEELIK | -0.0139 | 0.00984 | -1.41 | 0.159 | 1018 | 5;1013 |
| rs6733839 | AAPQWCQGK | -0.0378 | 0.0139 | -2.72 | 0.00674 | 1015 | 5;1010 |
| rs6733839 | AEEELIK | 0.0393 | 0.0558 | 0.705 | 0.481 | 1018 | 5;1013 |
| rs6733839 | AQPSDNAPAK | 0.0164 | 0.0407 | 0.402 | 0.688 | 1012 | 5;1007 |
| rs6733839 | VNHEPEPAGGATPGATLPK | 0.0281 | 0.0292 | 0.964 | 0.336 | 1017 | 5;1012 |
| rs6733839 | GPPVPPPPK | -0.00985 | 0.0125 | -0.788 | 0.431 | 1018 | 5;1013 |
| rs7274581 | LQAHLVAQTNLLR | 0.00807 | 0.0177 | 0.456 | 0.648 | 1018 | 5;1013 |
| rs7274581 | NQAEEELIK | 0.0132 | 0.0139 | 0.948 | 0.344 | 1018 | 5;1013 |
| rs7274581 | AAPQWCQGK | 0.00147 | 0.0197 | 0.0743 | 0.941 | 1015 | 5;1010 |
| rs7274581 | AEEELIK | -0.111 | 0.0786 | -1.41 | 0.159 | 1018 | 5;1013 |
| rs7274581 | AQPSDNAPAK | -0.0721 | 0.0575 | -1.25 | 0.21 | 1012 | 5;1007 |
| rs7274581 | VNHEPEPAGGATPGATLPK | -0.0748 | 0.0413 | -1.81 | 0.0702 | 1017 | 5;1012 |
| rs7274581 | GPPVPPPPK | -0.0151 | 0.0176 | -0.855 | 0.393 | 1018 | 5;1013 |
| rs9271192 | LQAHLVAQTNLLR | 0.0159 | 0.0107 | 1.48 | 0.139 | 1018 | 5;1013 |
| rs9271192 | NQAEEELIK | 0.00598 | 0.00844 | 0.709 | 0.478 | 1018 | 5;1013 |
| rs9271192 | AAPQWCQGK | 0.0116 | 0.012 | 0.967 | 0.334 | 1015 | 5;1010 |
| rs9271192 | AEEELIK | -0.041 | 0.0478 | -0.859 | 0.39 | 1018 | 5;1013 |
| rs9271192 | AQPSDNAPAK | 0.0389 | 0.0347 | 1.12 | 0.263 | 1012 | 5;1007 |
| rs9271192 | VNHEPEPAGGATPGATLPK | -0.0246 | 0.025 | -0.984 | 0.325 | 1017 | 5;1012 |
| rs9271192 | GPPVPPPPK | 0.00847 | 0.0107 | 0.791 | 0.429 | 1018 | 5;1013 |
| rs9331896 | LQAHLVAQTNLLR | -0.0196 | 0.0101 | -1.94 | 0.0529 | 1018 | 5;1013 |
| rs9331896 | NQAEEELIK | -0.00154 | 0.00798 | -0.193 | 0.847 | 1018 | 5;1013 |
| rs9331896 | AAPQWCQGK | 0.00139 | 0.0113 | 0.123 | 0.902 | 1015 | 5;1010 |
| rs9331896 | AEEELIK | 0.0425 | 0.0452 | 0.94 | 0.347 | 1018 | 5;1013 |
| rs9331896 | AQPSDNAPAK | 0.115 | 0.0327 | 3.51 | 0.000475 | 1012 | 5;1007 |
| rs9331896 | VNHEPEPAGGATPGATLPK | 0.0252 | 0.0237 | 1.06 | 0.288 | 1017 | 5;1012 |
| rs9331896 | GPPVPPPPK | 0.00417 | 0.0101 | 0.412 | 0.681 | 1018 | 5;1013 |
| rs983392 | LQAHLVAQTNLLR | 0.0124 | 0.0097 | 1.28 | 0.201 | 1018 | 5;1013 |
| rs983392 | NQAEEELIK | 0.00558 | 0.00764 | 0.731 | 0.465 | 1018 | 5;1013 |
| rs983392 | AAPQWCQGK | 0.0115 | 0.0108 | 1.06 | 0.287 | 1015 | 5;1010 |
| rs983392 | AEEELIK | 0.0094 | 0.0432 | 0.217 | 0.828 | 1018 | 5;1013 |
| rs983392 | AQPSDNAPAK | -0.00949 | 0.0314 | -0.302 | 0.763 | 1012 | 5;1007 |
| rs983392 | VNHEPEPAGGATPGATLPK | -0.000508 | 0.0227 | -0.0224 | 0.982 | 1017 | 5;1012 |
| rs983392 | GPPVPPPPK | -0.00196 | 0.00969 | -0.202 | 0.84 | 1018 | 5;1013 |

Suppl. Table 11

| outcome | Peptides | beta | se | tstat | pval | n | fdr |
|---|---|---|---|---|---|---|---|
| tangles_sqrt | LQAHLVAQTNLLR | -0.8 | 0.243 | -3.29 | 0.0011 | 430 | 0.0077 |
| tangles_sqrt | NQAEEELIK | -0.495 | 0.316 | -1.57 | 0.118 | 430 | 0.34533333 |
| tangles_sqrt | AAPQWCQGK | -0.33 | 0.227 | -1.45 | 0.148 | 430 | 0.34533333 |
| tangles_sqrt | AEEELIK | 0.00454 | 0.0574 | 0.0791 | 0.937 | 430 | 0.937 |
| tangles_sqrt | AQPSDNAPAK | -0.051 | 0.0799 | -0.639 | 0.523 | 430 | 0.91525 |
| tangles_sqrt | VNHEPEPAGGATPGATLPK | -0.0294 | 0.108 | -0.271 | 0.786 | 430 | 0.917 |
| tangles_sqrt | GPPVPPPPK | -0.128 | 0.306 | -0.419 | 0.675 | 430 | 0.917 |

Suppl. Table 12

| **Peptides** | **covariates** | **beta** | **se** | **tstat** | **pval** | **n** | **fdr** |
|---|---|---|---|---|---|---|---|
| LQAHLVAQTNLLR | age_death;msex | -0.000237 | 0.00021 | -1.13 | 0.261 | 579 | 0.7175 |
| NQAEEELIK | age_death;msex | -0.000182 | 0.000156 | -1.17 | 0.242 | 579 | 0.7175 |
| AAPQWCQGK | age_death;msex | -0.000325 | 0.000181 | -1.8 | 0.0725 | 579 | 0.7175 |
| AEEELIK | age_death;msex | 0.000753 | 0.000914 | 0.824 | 0.41 | 579 | 0.7175 |
| AQPSDNAPAK | age_death;msex | 0.000249 | 0.00068 | 0.367 | 0.714 | 578 | 0.768923077 |
| VNHEPEPAGGATPGATLPK | age_death;msex | 0.000208 | 0.000474 | 0.438 | 0.661 | 578 | 0.768923077 |
| GPPVPPPPK | age_death;msex | -0.000201 | 0.000162 | -1.24 | 0.215 | 579 | 0.7175 |
| LQAHLVAQTNLLR | age_death;msex;tangles_sqrt;amyloid_sqrt | -5.45E-05 | 0.00021 | -0.26 | 0.795 | 579 | 0.795 |
| NQAEEELIK | age_death;msex;tangles_sqrt;amyloid_sqrt | -0.000107 | 0.000156 | -0.683 | 0.495 | 579 | 0.768923077 |
| AAPQWCQGK | age_death;msex;tangles_sqrt;amyloid_sqrt | -0.000261 | 0.000184 | -1.42 | 0.155 | 579 | 0.7175 |
| AEEELIK | age_death;msex;tangles_sqrt;amyloid_sqrt | 0.000786 | 0.000934 | 0.842 | 0.4 | 579 | 0.7175 |
| AQPSDNAPAK | age_death;msex;tangles_sqrt;amyloid_sqrt | 0.000271 | 0.000694 | 0.39 | 0.697 | 578 | 0.768923077 |
| VNHEPEPAGGATPGATLPK | age_death;msex;tangles_sqrt;amyloid_sqrt | 0.00026 | 0.000484 | 0.537 | 0.592 | 578 | 0.768923077 |
| GPPVPPPPK | age_death;msex;tangles_sqrt;amyloid_sqrt | -0.000164 | 0.000166 | -0.988 | 0.324 | 579 | 0.7175 |
